# A molecular stabiliser of an inhibitory eIF2B-eIF2(αP) complex activates the Integrated Stress Response

**DOI:** 10.1101/2025.09.25.678332

**Authors:** Fiona Shilliday, Miguel Gancedo-Rodrigo, Ginto George, Santosh Adhikari, Syedah Neha Ashraf, Evelyne J. C. Barrey, Paolo A. Centrella, Damian Crowther, Paige Dickson, Diana Gikunju, Marie-Aude Guié, John P. Guilinger, Anders Gunnarsson, Heather P. Harding, Christopher D. Hupp, Rachael Jetson, Anthony D. Keefe, JeeSoo Monica Kim, Richard J. Lewis, Taiana Maia de Oliveira, Jennifer Le-Marshall, Usha Narayanan, Katherine A. Nugai, Dušan Petrović, Emma Rivers, David Ron, Daisy Stringfellow, Karl Syson, Lewis Ward, John T. S. Yeoman, Yan Yu, Ying Zhang, Alisa Zyryanova, David Baker, Perla Breccia, John Linley

**Affiliations:** Discovery Sciences, R&D, AstraZeneca, Cambridge, UK; Cambridge Institute for Medical Research (CIMR), University of Cambridge, Cambridge, UK; X-Chem Inc., Waltham, Massachusetts, USA; Discovery Sciences, R&D, AstraZeneca, Gothenburg, Sweden; Neuroscience, R&D, AstraZeneca, Cambridge, UK; Department of Medicinal Chemistry, Research and Early Development, Respiratory and Immunology (R&I), BioPharmaceuticals R&D, AstraZeneca, Gothenburg, Sweden; Biologics Engineering, R&D, AstraZeneca, Cambridge, UK

## Abstract

Eukaryotic initiation factor 2B (eIF2B), a guanine nucleotide exchange factor (GEF), promotes protein synthesis by charging translation initiation factor 2 (eIF2) with GTP. Stress-induced phosphorylation of eIF2 on its α-subunit [eIF2(αP)] inhibits this reaction triggering a protective Integrated Stress Response (ISR). A DNA-encoded chemical library (DEL) screen for modulators of eIF2B, led to the identification of a chemical series that inactivates eIF2B, stimulating the ISR. Cryo-EM of compound-bound eIF2B revealed a conformational switch to the inactive state engaged by eIF2(αP). In cells, compound activity was sensitive to eIF2’s phosphorylation state and to a competing eIF2B ligand (ISRIB) that activates the GEF allosterically. These findings mark the discovery of a first-in-class drug-like allosteric inhibitor of eIF2B, an ISR activator (ISRAC), paving the way to explore the therapeutic potential of eIF2B-directed ISR activation.

## Main

The preservation of eukaryotic cell homeostasis upon diverse cellular stress stimuli such as protein misfolding, nutrient deprivation, viral infection, oxidative stress or environmental conditions is mediated by the activation of a conserved cytoprotective signalling mechanism called the Integrated Stress Response (ISR)^1^. This self-defensive pathway is activated by stress-sensing kinases (PERK, PKR, HRI and GCN2) which specifically phosphorylate serine 51 of the α-subunit of the heterotrimeric GTPase, eukaryotic translation initiation factor 2 (eIF2), situated in its N-terminal domain (NTD)^2^. Guanine nucleotide exchange factor (GEF) eIF2B activates eIF2 by exchanging GDP to GTP at the *γ*-subunit allowing eIF2 to associate with methionyl initiator tRNA to form the eIF2-GTP-Met-tRNA_i_ ternary complex implicated in efficient mRNA translation initiation^3^. Phosphorylated eIF2 [eIF2(αP)] is an inhibitory allosteric regulator of eIF2B’s GEF activity. The resulting decline in ternary complexes attenuates translation of most mRNA, lowering global protein synthesis rates. Translation of a small set of stress-responsive mRNAs, exemplified by the transcription activator *ATF4*, that is basally-repressed in homeostatic conditions, is activated by eIF2(αP) in a mechanism dependent on upstream open reading frames (uORFs)^4,5^. ISR-activated reprogramming of translation and gene expression contributes to recovery from stress but if sustained can promote apoptosis^6^.

Failure to induce or downregulate ISR signalling is linked to the pathogenesis of cancer, neurologic and metabolic disorders^6^. Best documented are loss-of-function mutations in genes encoding ISR components that directly or indirectly compromise eIF2’s phosphorylated state and ternary complex levels. Mutations affecting eIF2 and eIF2B which activate the ISR are well-characterised, as reviewed in Hanson et al. 2022^7^.

eIF2B is formed from two tetramers of beta, gamma, delta and epsilon subunits (eIF2Bβγδε). These come together to form an octamer with regulatory β and δ subunits being at the core and catalytic γ and ε at the exterior. A boost in stability occurs when a dimer of eIF2B alpha (eIF2Bα) joins the octamer to form a decamer with a 20-fold enhancement of eIF2B’s nucleotide exchange activity^8^. eIF2B’s substrate, unphosphorylated eIF2, and its allosteric-inhibitory phosphorylated form, eIF2(αP), bind differently. These independent binding events favor, and in turn stabilize, two structurally distinct conformations of eIF2B: an active A-state and an inactive I-state^8–11^. In the A-state, eIF2B carries out its GEF activity. Its core subunits adopt an inward conformation, allowing eIF2 to bind as a substrate and exchange its GDP for GTP. Conversely, the I-state exhibits an outward conformation favouring eIF2(αP) binding as an inhibitor^10,11^.

Pharmacological modulation of the ISR targeting the central eIF2B-eIF2 hub was first realised by the discovery of ISRIB^12^, a drug-like small molecule which directly binds and stabilises the eIF2B symmetric core imposing the A-state to allosterically antagonise eIF2(αP)’s inhibitory effect^8–11,13,14^. ISRIB’s beneficial properties in reversing the ISR have been shown in neurodegeneration and brain injury models, followed by ISRIB-like analogue 2BAct^15^ and more recently DNL-343^16^.

Promotion of the A-state, favoured by ISRIB binding to an allosteric regulatory site of eIF2B, is a well-established mechanism of action (MOA) of an ISR inhibitory compound. Differences in structure of the allosteric site between the A and I-state of eIF2B preclude ISRIB binding to the latter and suggested the possibility of modulating eIF2B in the opposite direction, through compounds that bind to and stabilize the I-state, thus activating the ISR^10,11^. This notion is further supported by the recent description of a single point mutation pushing eIF2B towards the I-state^17^: eIF2B β-subunit H160 is situated in the ISRIB binding pocket. Overlay of wild-type eIF2B and eIF2BβH160D shows widening of the pocket in the mutant - a change that propagates across the decamer partially inducing the I-state of eIF2B - raising the spectre of compounds achieving a similar outcome.

Here, we used DNA-encoded chemical library (DEL) screening and medicinal chemistry, combined with biochemical, biophysical, structural, and cellular analyses to discover and characterise a first-in-class drug-like molecule targeting eIF2B and activating the ISR pathway, an ISRAC (Integrated Stress Response Activator).

## Results

### Identification of Compound A, a small molecule binding to eIF2B in complex with phosphorylated eIF2

To identify small-molecule modulators of eIF2B, we screened a mixture of 26 DNA-encoded chemical libraries containing 1.55 billion different compounds for eIF2B binders under a range of conditions in parallel (Fig. 1A). As a counter-screen, biotinylated eIF2α and (P)eIF2α were included in separate conditions to identify compounds which interact with eIF2B binding partners.

**Fig. 1:**
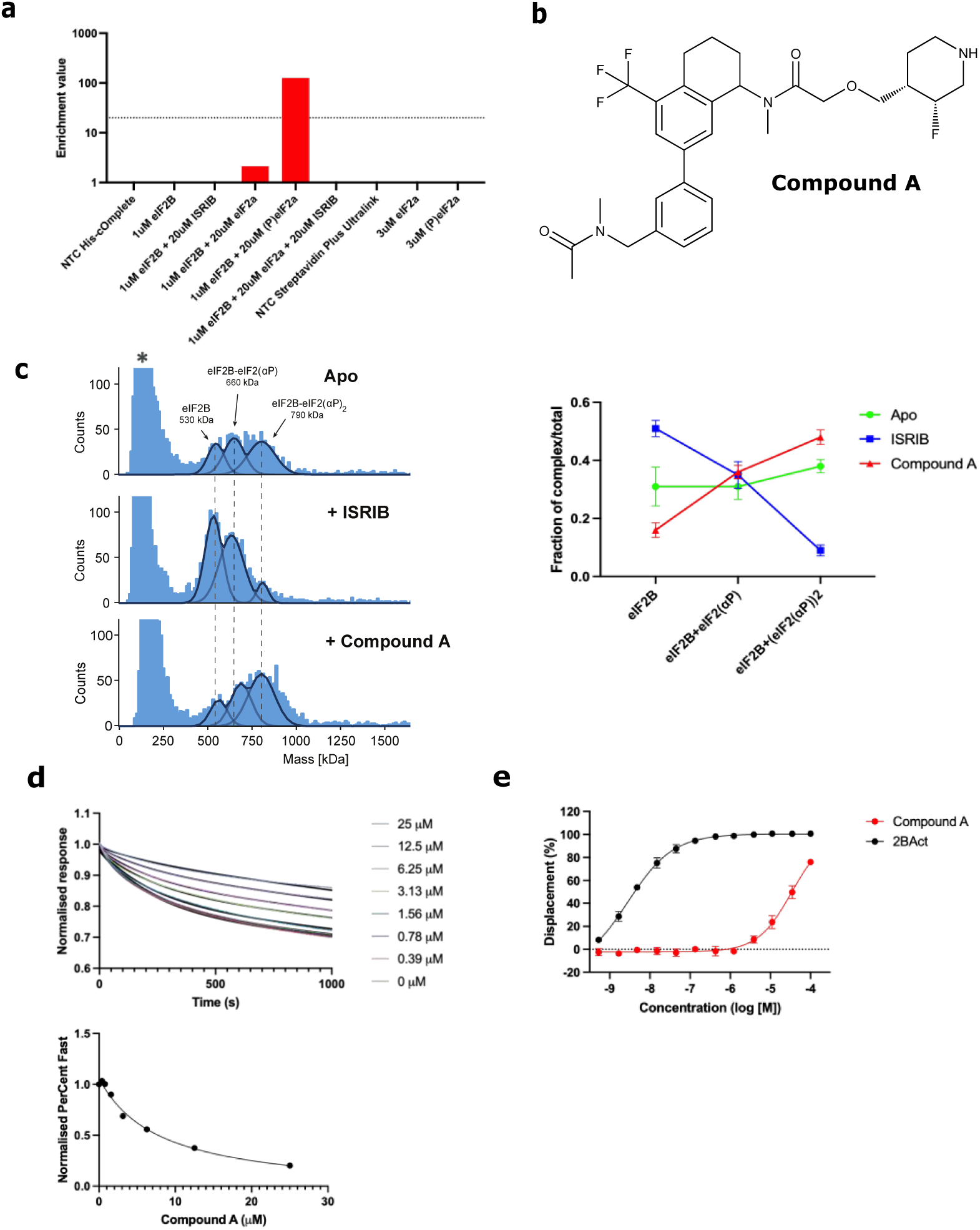
DEL-enabled discovery of a compound that stabilises the inhibitory eIF2B:eIF2(aP) complex. **a,** DEL screening conditions used in parallel for the identification of eIF2B modulators. No-target control (1, IMAC matrix), eIF2B alone (2), eIF2B bound to ISRIB (3), eIF2B in complex with eIF2a (4), eIF2B in complex with (P)eIF2a (5) and the latter in the presence of ISRIB (6), No-target control (7, Streptavidin matrix), biotinylated eIF2a (8) and (P)eIF2a (9) DNA-tagged ISRIB was included as an Internal Positive Control and a corresponding DNA-tagged linker was included as an Internal Negative Control (Extended Data Fig. 1). Hits were grouped into clusters and Series 1 was only observed in condition (5) with a statistically significant enrichment value of >20 (indicated by the dotted line). Compound A was prioritised for off-DNA synthesis and progressed for further characterisation. **b**, The chemical structure of Compound A. **c**, Histogram of the size distribution of complexes formed between eIF2B and eIF2(aP), in absence of compound (Apo) and in presence of ISRIB or Compound A, as revealed by mass photometry. The three separate peaks are labelled with their corresponding expected MWs. Note, the eIF2B sample also contains lower molecular weight species with a mass consistent with individual dissociated subunits and dimers thereof (marked *) (left panel). The fraction of each of the three complexes in each sample is quantified (right panel). **d**, Upper panel: Time-dependent change to the interferometric signal from dissociation of eIF2B from immobilised (P)eIF2a-NTD on a BLI sensor in presence of the indicated concentrations of Compound A. The signal, in nm, was normalised to the initial values in each trace, corresponding to the amount of biotinylated (P)eIF2a-NTD:eIF2B formed on the sensor. Lower panel: The fraction of the dissociation attributed to the fast phase (PercentFast) calculated from the biphasic decay fits shown on the upper panel was plotted as a function of the concentration of Compound A. Values were normalised to the PercentFast value of the dissociation recorded in absence of compound. Compound A decreases the Percent Fast by ≥ 82% with an EC50 of 7.9 mM (4.3 - 16.7, 95% CI). Shown is one example of an n=3 experiment. **e**, Concentration-dependent changes to the binding of labelled ISRIB to eIF2B (as revealed by HTRF) in presence of the ISRIB analogue 2BAct or Compound A. Note the weak displacement of the ISRIB probe from eIF2B by Compound A (76% Activity at 100 mM).

The screening conditions were designed to inform the MOA of hits by establishing whether binding is competed by ISRIB, and whether hits preferentially engage the A-state eIF2B:eIF2〈 complex, the I-state eIF2B:(P)eIF2α complex or apo-eIF2B. DNA-tagged ISRIB was included as an internal positive control (Extended Data Fig. 1) and a fold-enrichment value over a DNA-tagged negative control of >20,000 was interpreted as highly statistically significant binding thereby validating the screening conditions.

Hits were identified across multiple selection conditions and grouped into clusters of structurally related compounds. **Series 1** was enriched when eIF2B was in complex with (P)eIF2α, but not significantly when eIF2B was complexed with non-phosphorylated eIF2α and not at all with eIF2B alone (Fig. 1A). Furthermore, the enrichment of **Series 1** was eliminated by the presence of ISRIB. Compound A (Fig. 1B), a mixture of two diastereomers, from **Series 1** was prioritised for off-DNA synthesis.

### Compound A stabilises the eIF2B decamer and favours eIF2B:eIF2(αP) complex formation

To explore the MOA for Compound A we first examined its effect using interferometric scattering microscopy (ISCAT), also known as mass photometry. In samples of recombinantly expressed eIF2B, ISCAT confirmed the presence of the decameric eIF2B complex (αβγδε)_2_ at ∼530 kDa as well as a broad peak consistent in mass with individual subunits or dimers (Extended Data Fig. 2A). ISCAT of eIF2 and eIF2(αP) confirmed homogenous populations of the expected mass (Extended Data Fig. 2B and C). Upon dilution of eIF2B to concentrations used for ISCAT measurements, the complex slowly dissociated on a timeframe of hours. Differences in decamer dissociation in the absence or presence of compound confirmed the previously observed stabilization of eIF2B by ISRIB^8^ (Extended Data Fig. 2D and E). Compound A behaved in a similar manner. Upon mixing of eIF2B with either eIF2 or eIF2(αP) three complexes with different masses were observed: i) eIF2B alone, ii) eIF2B with one eIF2/ eIF2(αP) and iii) eIF2B-with two eIF2/ eIF2(αP). By quantifying the relative abundance of these complexes, it was possible to deduce mechanistic insights that differentiated the two compounds from Apo as well as from each other. ISRIB decreased the affinity of eIF2B for eIF2(αP) resulting in less eIF2B-(eIF2(αP))_2_ complex (compared to the apo-eIF2B). Compound A acted in the opposite manner, stabilising the eIF2B-eIF2(αP) interaction resulting in more eIF2B-(eIF2(αP))_2_ complex (Fig. 1C). Interestingly, the compounds had no detectable effect on the corresponding complexes involving unphosphorylated eIF2 (Extended Data Fig. 2F).

The isolated phosphorylated N-terminal domain of eIF2a, (P)eIF2α-NTD, recapitulates features of the phosphorylated trimer^9,10^. Therefore we monitored Compound A’s effect on the complex formed between biotinylated (P)eIF2α-NTD immobilised on a Biolayer Interferometry (BLI) sensor and eIF2B (modified from Zyryanova et al, 2021^10^). The dissociation phase of a preformed complex was measured in absence or presence of increasing concentrations of Compound A (Fig. 1D). As previously reported^10^, in absence of compound both the association and dissociation phases were multi-phasic, likely reflecting multiple binding events that comprise this interaction. Compound A reduces the dissociation rate of the eIF2B:(P)eIF2α-NTD complex in a concentration-dependent manner. To quantify this effect, we fitted the dissociation traces to a two-phase model, which describes the sum of a fast and a slow first order decay. We found that Compound A decreased the contribution of the fast component to the overall dissociation (Percent Fast) by 82% with an EC_50_ of 7.9μM. This is the opposite effect of ISRIB, which increases the dissociation rate of eIF2B from the immobilised (P)eIF2α−NTD.

In a Homogeneous Time-Resolved Fluorescence (HTRF) assay that tracks displacement of an ISRIB probe from eIF2B, Compound A was confirmed as a hit but only showed weak displacement of ISRIB (76% Activity at 100 μM, Fig. 1E). This was consistent with the DEL screening output, in which Compound A was not enriched in the presence of eIF2B and saturating concentrations of eIF2 α or in the presence of ISRIB. It also suggested that Compound A could bind eIF2B alone, but at levels below the detection limit of the DEL screen.

### Compound A-*(S)* is the more active stereoisomer showing cooperative binding to eIF2B and reducing GEF activity

Compound A has two diastereoisomeric forms generated by the stereochemistry (*S* or *R*) at the tetrahydronaphthalene carbon 7 (Fig. 2A), with the two additional stereocentres on the piperidine group being fixed as (3*R*,4*S*)-3-fluoropiperidine. To assess their potential significance, we compared the effect of each isolated diastereoisomer on stability of the complex formed between immobilized (P)eIF2α-NTD and eIF2B in the BLI assay described above. Compound A-(*S*) refers to where stereochemistry at the tetrahydronaphthalene carbon 7 is defined as *S* and Compound A-(*R*) refers to where stereochemistry at the tetrahydronaphthalene carbon 7 is defined as *R*.

**Fig. 2:**
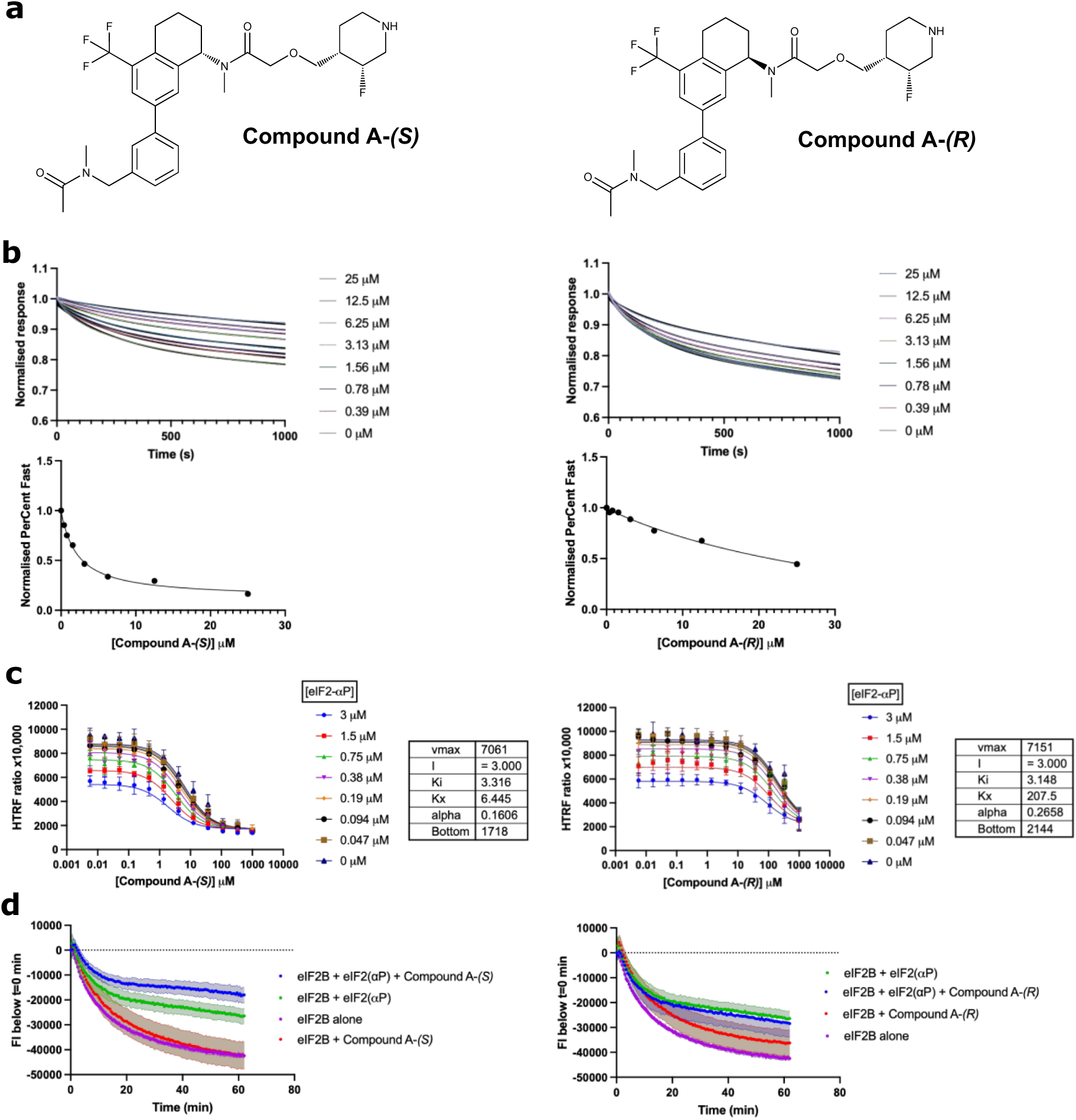
Stereo-specificity in Compound A: The *(S)* stereoisomer is more active than the *(R)* stereoisomer. **a,** Structures of Compound A -*(S)* and -*(R)* stereoisomers. **b,** Upper panels: Time dependent change to the interferometric signal from dissociation of the eIF2B:(P)eIF2a-NTD complex immobilised on a BLI sensor in presence of the indicated concentrations of Compound A-*(S)* and *(R).* Lower panels: The fraction of the dissociation attributed to the fast phase (PercentFast) calculated from the biphasic decay fits shown on the upper panel was plotted as a function of the concentration of Compound A-*(S)* and *(R)* (as in Fig. 1d). Compound A-*(S)* (top) has a greater stabilisation effect of the eIF2B:(P)eIF2a-NTD interaction, decreasing the Percent Fast value by 82% with an EC50 of 2.2 mM (1.5 - 3.2, 95% CI), than Compound A-*(R)* (bottom), which decreased the Percent Fast value by 49% with an EC50 of 39.2 mM (17.5 - 235.7, 95% CI). **c,** In HTRF matrix experiments the *S* stereoisomer displayed ∼30-fold greater potency over the *R* stereoisomer, with apparent K_½ max_ values of 6.4 and 207.5 mM respectively. From the Yonetani-Theorell fitting the alpha value which indicates that both stereoisomers have cooperative binding with eIF2(aP), with values of 0.16 (*S*) and 0.27 (*R*). **d,** Traces of time-dependent fluorescent intensity arising from BODIPY-GDP loaded eIF2 in presence of eIF2B alone (purple), eIF2B and eIF2(aP) (green), eIF2B and Compound A (red), eIF2B, Compound A and eIF2(aP) (blue). The *S* and *R* stereoisomers are shown in the upper and lower panels. Decline in fluorescent intensity reflects the GEF activity of eIF2B. Both stereoisomers reduce the GEF activity of eIF2B with Compound A*-(S)* having a two-fold longer half-life than Compound A-*(R*), (76.42 s (69.98 - 83.88, 95% CI vs 33.65 s (31.45 - 36.03, 95% CI).

Compound A-(*S*) was found to be the more active of the two stereoisomers, decreasing the Percent Fast value by 82% with an EC_50_ of 2.2 μM whilst Compound A-(*R*) decreased the Percent Fast value by 49% with an EC_50_ of 39.2 μM (Fig. 2B).

To further probe the nature of the binding between Compound A stereoisomers and (P)eIF2α, matrix experiments were performed by HTRF. Compound A-(*S*), Compound A-(*R*) and (P)eIF2α could all displace the ISRIB probe (Fig. 2C). The *S* stereoisomer displayed ∼30-fold greater potency over the *R* stereoisomer, with K_½ max_ values of 6.4 and 207.5 μM respectively. The K_½ max_ for (P)eIF2α remained constant at 3 μM, with both the *R* and *S* stereoisomer. The alpha value from Yonetani-Theorell fitting, gives a measure of cooperativity in ligand binding, α = 1 indicates that each ligand binds independently, α = > 1 indicates ligands have negative cooperativity and α < 1 indicates positive cooperativity. Interestingly, both stereoisomers showed cooperative binding with (P)eIF2〈, with alpha values of 0.16 and 0.27 for the *S* and *R* stereoisomers respectively.

To investigate the functional consequences of Compound A stereoisomers binding on eIF2B’s GEF activity towards GDP-loaded eIF2 we measured the rate of GDP-BODIPY dissociation from eIF2 in absence and presence of compound [reflected in a decrease in fluorescence intensity (FI)]. Both stereoisomers attenuated GEF activity, in this assay too the *S* stereoisomer was more active imparting a greater than two-fold longer half-life (t1/2) on the eIF2B-bound GDP-BODIPY compared to the *R* stereoisomer [76.42 (69.98-83.88 95% CI) vs 33.65 (31.45-36.03, 95% CI)]. Stabilisation of the bound GDP-BODIPY by either stereoisomer was eIF2(αP)-dependent, consistent with the observations in the HTRF assay (Fig. 2D).

### Compound A-(*S*) binds the ISRIB pocket of eIF2B, favouring the I-state

Compound A was only enriched in the DEL screen in presence of both eIF2B and eIF2(αP). Therefore, we proposed to determine the structure of eIF2B+(P)eIF2α-NTD+Compound-A-*(S)* to explore the compound’s MOA and reveal its binding site.

A cryo-EM structure of eIF2B+(P)eIF2α-NTD+Compound-A-*(S)* was determined to an overall resolution of 2.8 Å. A density corresponding to Compound A-*(S)* was observed between the beta and delta subunits of eIF2B’s core at the ISRIB-binding site (Fig. 3A, Supplementary Table 1, PDB: ????, EMDB: ?????). An overlay with eIF2B+eIF2(〈P) structure^9^ (PDB 6K72) showed good overall agreement in the positioning of all subunits, notably in the disposition of the phosphorylated eIF2α subunit’s N-terminal domain that is docked between eIF2Bα, eIF2Bβ and eIF2B™, a binding pose correlated with inhibition of eIF2B’s GEF activity^8,9^ (Fig. 3A). These features, suggestive of eIF2B’s I-state, distinguish the ternary complex of eIF2B + (P)eIF2α-NTD + Compound A-*(S)* from all known ternary complexes of eIF2B with ISRIB, which are only recovered with non-phosphorylated eIF2a and are in the A-state^10^.

**Fig. 3:**
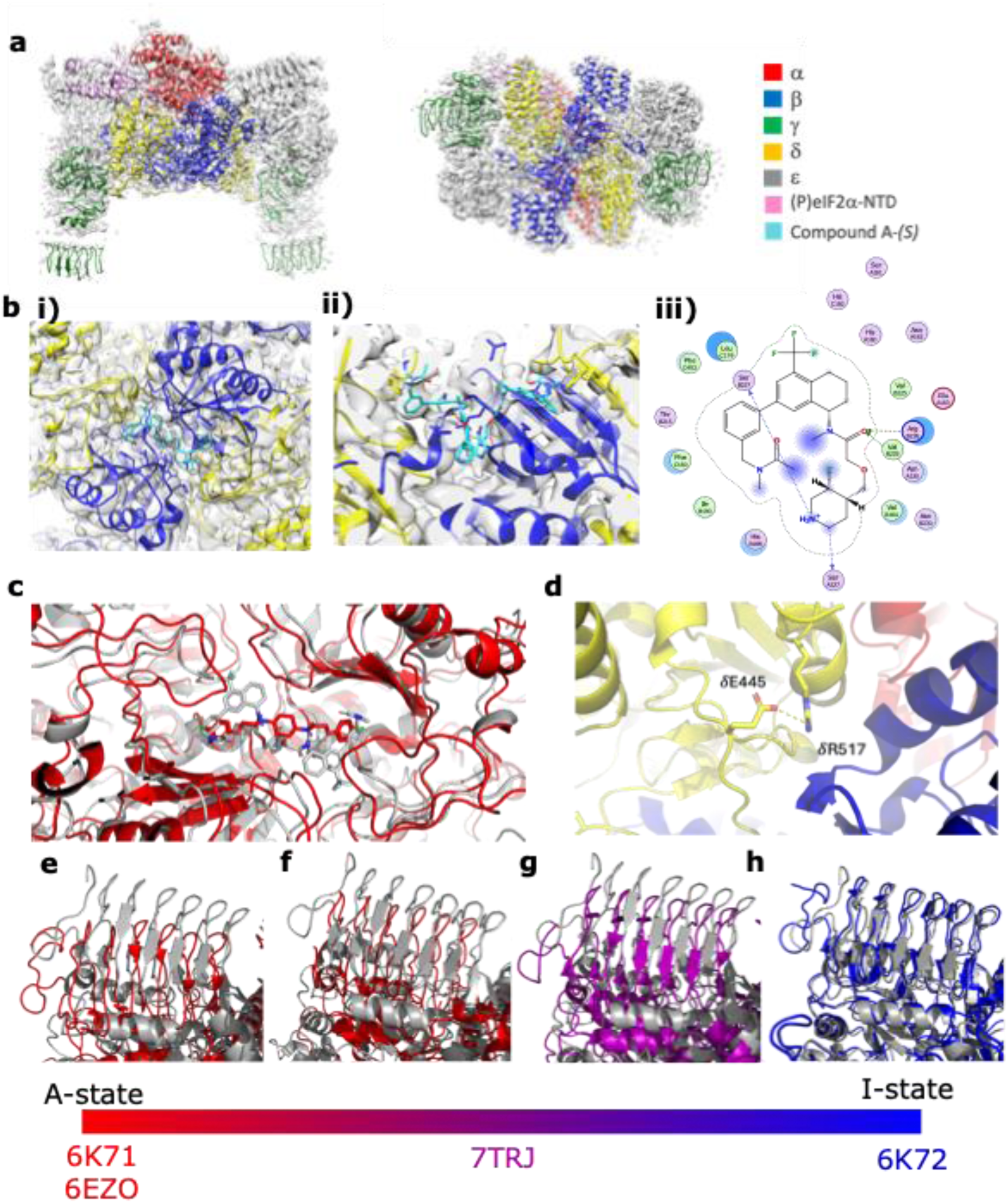
Cryo-EM structure of Compound A-*(S)* bound to the eIF2B:PeIF2a complex. **a,** Cryo-EM map of an eIF2B: PeIF2a-NTD complex with bound Compound A-*(S)* (transparent grey) (EMDB: ?????) and model (PDB: ????) with each eIF2B subunit and the PeIF2a-NTD coloured as listed in the key. Note the position of (P)eIF2a-NTD between the eIF2Ba, eIF2Bb and eIF2Bd, a binding poise correlated with eIF2B’s inhibitory I-state. **b,** Panel *i*-inset view of the eIF2B beta and delta subunit core showing additional density which accommodates a complete copy of Compound A-*(S)* on one side and the headgroup of a second molecule in the opposing pentamer. Panel *ii*-inset view showing that only one piperidine tail can be positioned at once (right hand side compound shown positioned in density) whereas on the opposing side the piperidine tail cannot be accommodated so is pushed out of density (second molecule omitted completely in deposited model). Panel *iii*-Compound A-*(S)* and its surrounding residues in an interaction map generated by MOE. **c,** Overlay of eIF2B:(P)eIF2a-NTD:Compound-A-*(S)* structure (shown in grey) with 6EZO eIF2B + ISRIB structure (shown in red) aligned by their eIF2Ba subunits. Note the widening and lengthening of the pocket when Compound A-*(S)* is bound. **d,** inset view of alpha and delta subunits of eIF2B where there are two differing salt bridges formed when eIF2B is in the A or I-state. Here residues eIF2BdR517 and eIF2BdE445 are observed to form the I-state salt-bridge. **e, f, g and h**, overlays of eIF2B:(P)eIF2a-NTD:Compound-A-*(S)* structure (in grey) with the indicated structures (colour coded to their position on the A / I-state scale depicted below). – **e,** 6K71 eIF2B:eIF2 shows an angular shift of the epsilon subunits if the beta and delta core is aligned **f,** 6EZO eIF2B:ISRIB similarly to 6K71 showing the angular shift of the epsilon subunits. Both 6K71 and 6EZO are A-state structures so were not expected to overlay well with our structure. **g,** 7TRJ eIF2B with H160D mutation is representative of a partial I-state structure compared with the fully I-state eIF2B:(P)eIF2a:Compound-A-*(S)* structure. **h,** 6K72 eIF2B:eIF2(aP) shows good overall agreement as expected from an I-state structure.

Compound A-*(S)* was observed to interact with multiple eIF2B residues, including βR228 via a H-bond to the carbonyl of the tetrahydronaphthalenyl-acetamide in Compound A*-(S)* (Fig. 3B). The fluoro-piperidine tail was buried into the core of eIF2B with an H-bond from the nitrogen of the piperidine ring to backbone carbonyl of βS227 and another H-bond from the piperidine ring to S227in the opposing beta subunit across the dimer interface. The same tetrahydronaphthalenyl-acetamide carbonyl of Compound A-*(S)* also interacted with eIF2BβN230 and eIF2BβR228. In the opposite tetramer there was partial density for the trifluoromethyl-tetrahydronaphthalene core group but the fluoro-piperidine tail could not be positioned as its mirrored binding site was already occupied by the first molecule.

The trifluoromethyl-tetrahydronaphthalene core group of the molecule is positioned between eIF2B residues δL179, βE162 and βH188. Notably, all three of these residues also surround ISRIB in it’s bound form, making interactions with the compound^14^. While these residues do not form specific interactions with Compound A-*(S),* the position of them is altered compared to an ISRIB bound structure. Most prominently, δL179 is looking outwards from the centre of eIF2B, partially rationalising the opposite MOAs of Compound A-*(S)* and ISRIB, and the ability of Compound A-*(S)* to stabilise the interaction with eIF2(αP) and inhibit eIF2B’s GEF activity (Fig. 3C).

The cryo-EM map lacked density for the C-terminal tails of the eIF2B gamma subunits, suggestive of their flexibility (Extended Data Fig. 3B). We subjected the dataset to 3D variability analysis in cryoSPARC and showed the flexibility represented the oscillation between A and I-states in solution as previously observed by Schoof et al.^11^. A cluster of particles could also be isolated using this method that showed improved resolution for two copies of (P)eIF2α-NTD and the compound (Extended Data Fig. 3C and D).

Compound A-*(S)* binding is associated with widening and lengthening of the pocket between eIF2B’s beta and delta subunits, consistent with Compound A-*(S)’s* MOA to induce the I-state of eIF2B. An overlay with I-state structure eIF2B+eIF2(αP) (PDB 6K72) showed good agreement across all subunits whereas overlays with A-state structures of eIF2B+eIF2^9^ (PDB 6K71) and eIF2B+ISRIB^14^ (PDB 6EZO) showed a pronounced angular shift of the epsilon subunits when the core of eIF2B was aligned (Fig. 3E-H). There was density to support salt bridge formation between eIF2BδR517 and eIF2BδE445, as previously described by Lawrence et al.^18^ to be a characteristic of the I-state (Fig. 3D). The helix rotation and movement of δR517 was observed in an overlay with eIF2B+ISRIB (PDB 6CAJ) structure, and the A-state salt bridge between eIF2BδR517 and eIF2BαD298 was precluded. These conclusions are further supported by 3D variability analysis of an eIF2B+(P)eIF2α-NTD cryo-EM dataset obtained in the absence of any compound in which 50% of particles were in the I-state. Whereas in the presence of Compound A-(*S*) this was significantly increased to 95% of the particles.

### Compound A and Compound A-*(S)* modulate the ISR in cells

Compound A-*(S)* was shown to decrease GEF activity in vitro so, looking downstream in the ISR pathway we measured its ability to activate one of the ISR target genes, C/EBP homologous protein (CHOP). Compound A-*(S)* elicited a weak but reproducible activation of a CHOP::GFP reporter gene^19^ reaching a shallow plateau (Fig. 4A).

**Fig. 4:**
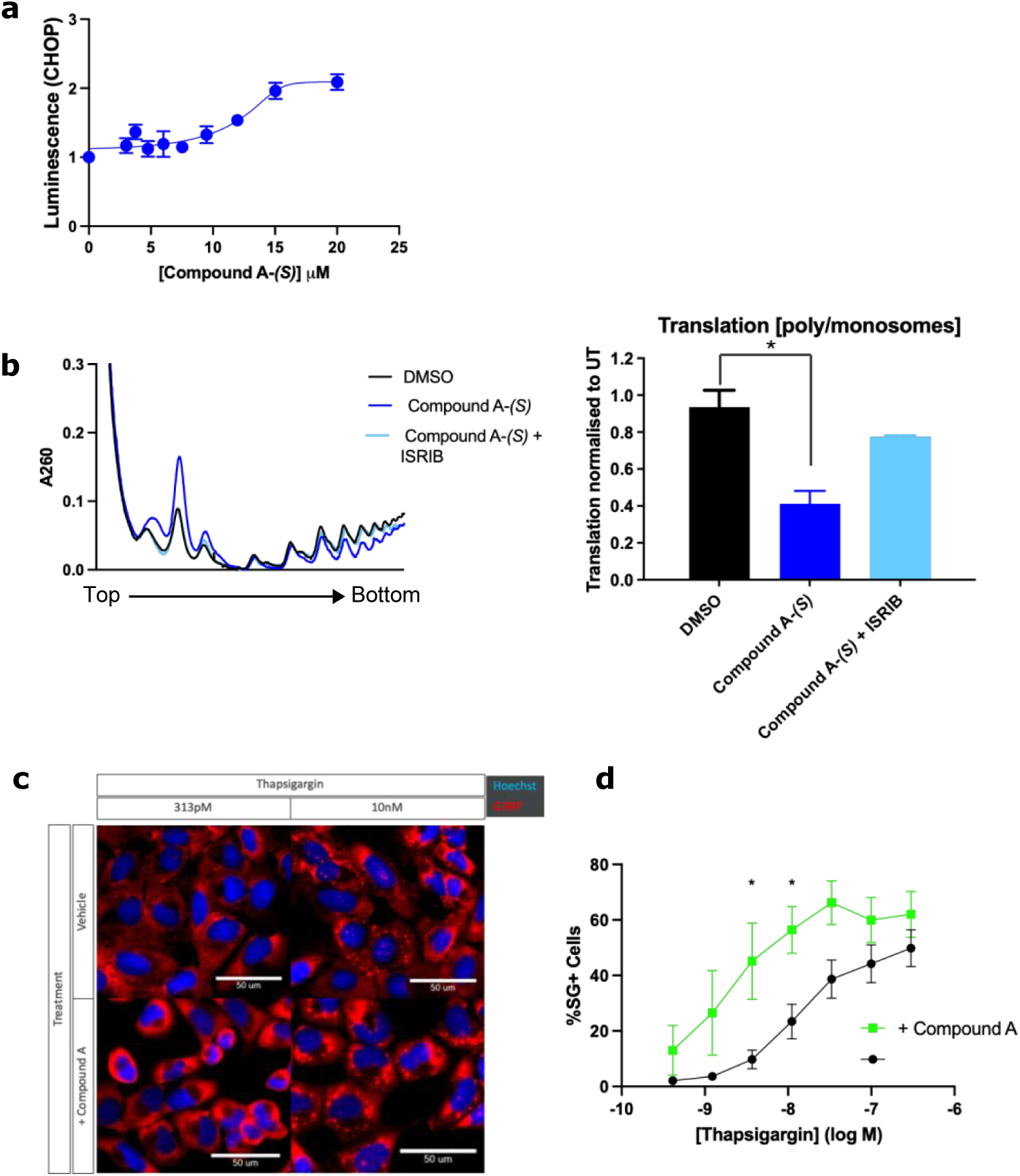
Cellular characterisation of Compound A and Compound A-*(S)* demonstrates modulation of the ISR. **a,** Activity of a CHOP::Luciferase ISR reporter gene plotted against the concentration of Compound A-*(S)* applied to CHO cells over 18 hours. Shown is the mean and SD, n = 3. The data were fitted by non-linear regression analysis to a dose response model using Prism GraphPad v10. EC_50_ 12.47 mM; (11.27-13.95 95% confidence intervals, CI). **b,** A260 traces from 10–60% sucrose gradients loaded with cytoplasmic extracts from cells treated with DMSO control (DMSO, black), Compound A-(*S*) (20 mM, purple), Compound A-(*S*) (20 mM) + ISRIB (2 mM, light blue). Quantification of the ratio of polysomes/ monosomes from three experiments. Statistical analysis was performed by a two-sided unpaired Welch’s t-test and significance is indicated by asterisks (* p<0.05). **c,** Representative fluorescent photomicrographs of G3BP (a marker of stress granule formation) and Hoechst 33258 (nuclear marker) in fixed U2OS cells exposed to the indicated concentration of thapsigargin in presence and absence of 30 mM Compound A. **d,** Quantification of data from experiments as in **c**, performed over a range of thapsigargin concentrations, 5-6 technical replicates per condition per biological run. * represents p < 0.05 with a 2way ANOVA multiple comparisons test with Sidak’s multiple comparisons test.

Engagement of eIF2B’s regulatory sites predicts inhibition of the initiation phase of translation, as cells are depleted of ternary complexes of eIF2, GTP and charged methionyl initiator tRNA whose formation depends on a steady supply of GTP-bound eIF2; the product of eIF2B’s enzymatic activity. This facet can be studied by polysome profiles, which separate actively translating ribosomes associated with mRNAs (polysomes) from inactive and dissociated ribosomal subunits. Fractionation of mammalian cell lysates based on their density (density across a sucrose gradient) revealed that untreated cells have more RNA (mainly derived from ribosomes) in dense fractions, corresponding to translationally active polysomes than cells treated with Compound A-*(S)*. Furthermore, the inhibitory effect of Compound A-*(S)* on translation initiation as reflected by the polysome profiles was reversed by ISRIB (Fig. 4B), consistent with competition between these two ligands for an over-lapping regulatory site on eIF2B, as suggested by the structure of Compound A-*(S)-*bound eIF2B and by the HTRF assay described above.

Consistent with inhibition of translation initiation and activation of an ISR target gene, Compound A also synergised with the ISR inducing agent thapsigargin (an activator of the eIF2 kinase PERK^20^) in promoting stress-granule formation (another marker of ISR activity): a clear leftward shift in the thapsigargin concentration response of stress granules was observed in cells exposed to Compound A (Fig. 4C & D).

These observations suggested that whilst Compound A had only weak ISR-inducing activity in cells its effects were nonetheless “on-pathway”, which motivated us to look for ISR activators based on the same mechanism as Compound A-*(S)’s* but with greater potency.

### Compound B: an ISR activator

Seeking stronger ISR activators, based on Compound A*-(S)*’s mode of eIF2B engagement in the cryo-EM structure, we synthesized a symmetric molecule containing two Compound A*-(S)* unique pharmacophores, predicted to position one in each of eIF2B’s two regulatory sites. A 1,2-dipropoxyethane linker was used to connect the two (trifluoromethyl)-tetrahydronaphthalenyl)acetamides while discarding the basic piperidine tail in the design. This appeared optimal in terms of predicted fit into the binding pocket although compromised the solubility of the resulting symmetric Compound B (Fig. 5A).

**Figure 5:**
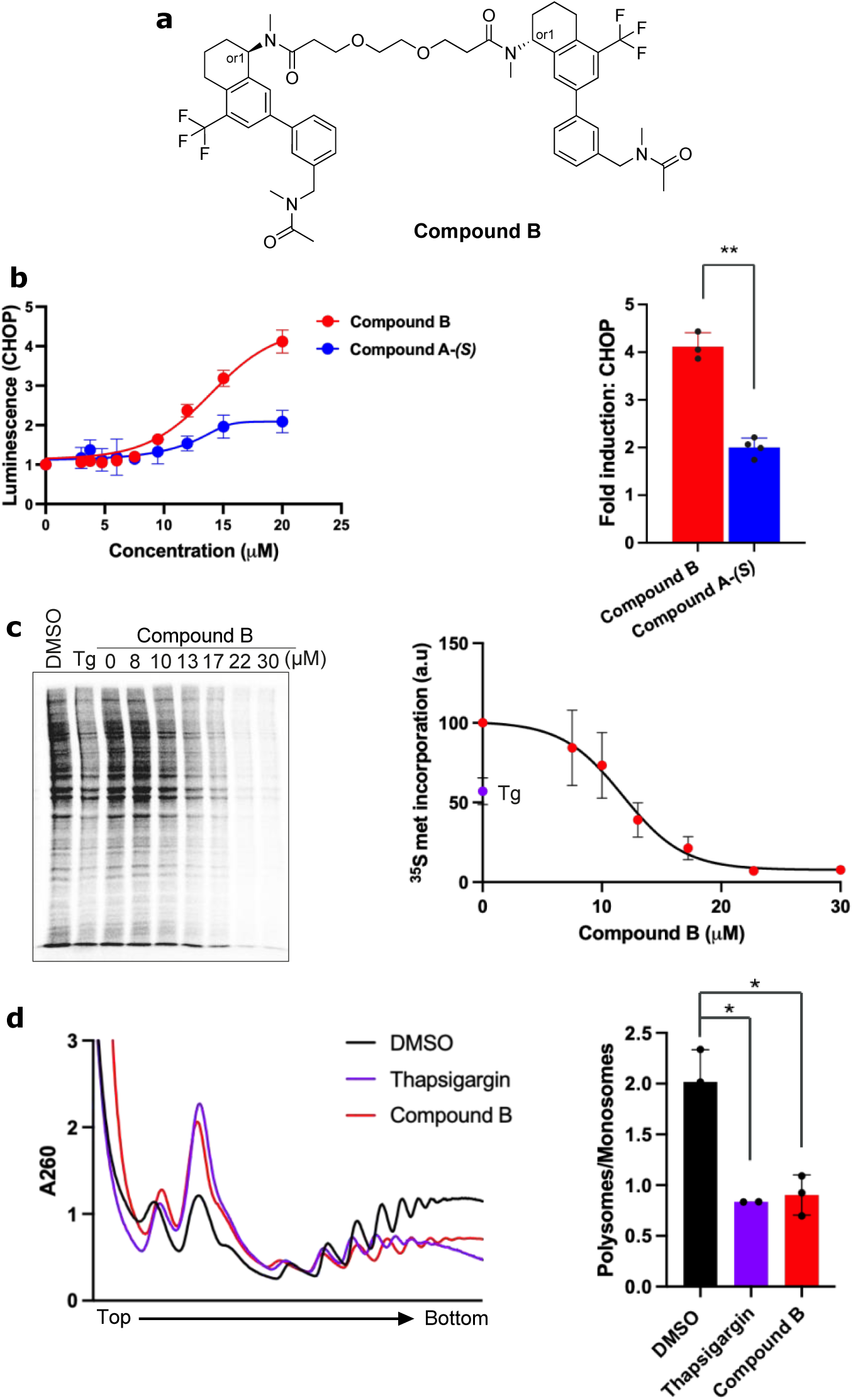
Compound B activates the ISR to a greater extent than Compound A-*(S)*. **a,** Chemical structure of Compound B. **b,** CHOP::Luciferase ISR reporter gene activity plotted against the concentration of Compound B applied to cells over 18 hours (the trace of Compound A-*(S),* from Figure 4A is provided as a reference). Shown is the mean and SD, n = 3. The data were fitted as in Figure 4A. The column graph to the right displays all four data points and the mean ± SD of the fold increase (over untreated) of samples exposed to 20 mM of the indicated compounds. Statistical analysis was performed by a two-sided unpaired Welch’s t-test and significance is indicated by asterisks (** p<0.01). **c,** Left panel: Autoradiograph of ^35^S methionine incorporation into newly synthesized cellular proteins over a 30 min period following exposure to saturating concentration of thapsigargin and to the indicated concentration of Compound B commencing 30 minutes before labelling and continued throughout the labelling period (representative of three such experiments). Right panel: Quantification of ^35^S methionine incorporation as a function of Compound B concentration (n = 3). The data were fitted by non-linear regression analysis to an inhibitor versus response model. IC_50_ = 11.74 mM (95% CI = 9.95 – 13.61). **d,** A260 traces from 10–60% sucrose gradients loaded with cytoplasmic extracts from cells treated with DMSO control (DMSO, black), thapsigargin (Tg, 1 mM, purple) or Compound B (30 mM, red), representative of three experiments. Quantification of the ratio of polysomes/ monosomes (n = 3). Statistical analysis was performed by a two-sided unpaired Welch’s t-test and significance is indicated by asterisks (* p<0.05).

The low solubility of Compound B in physiological buffers precluded assessment of its biochemical properties in vitro. Nonetheless, when applied to cultured cells Compound B reproducibly induced the CHOP::LUC ISR reporter gene with 2-fold greater plateau activation values than Compound A*-(S)* (Fig. 5B).

To generate a robust signal, the reporter gene-based assay requires many hours of exposure (favouring accumulation of reporter gene product). However, engagement of eIF2B in an inhibitory mode predicts rapid effects on protein synthesis rates, on the time scale observed in cells with mounting levels of phosphorylated eIF2. Therefore, we compared rates of ^35^S-methionine incorporation into newly synthesized proteins in cells exposed briefly (one hour) to a saturating concentration of thapsigargin or to escalating concentrations of Compound B. Compound B potently inhibited protein synthesis with an IC_50_ of 12 μM and plateau values nearing complete inhibition (Fig. 5C).

In both thapsigargin- and Compound B-treated cells, polysomes are depleted and the signal arising from monosomes increases (Fig. 5D). These observations are consistent with the hypothesised inhibition of eIF2B by phosphorylated eIF2 in thapsigargin-treated cells or by direct action in Compound B-treated cells.

Activation of the ISR was also observed in Compound B-treated cells bearing an integrated copy of CHOP::GFP, a different ISR transcriptional reporter. The EC_50_ value of Compound B action in the two assays is similar (13 μM (12.42-13.78 95% CI) in the CHOP::LUC assay, (Fig. 5B) and 15 μM (14.94-16.21 95% CI) using CHOP::GFP, Fig. 6A). ISRIB, for which the binding site and binding mode partially overlap with Compound B, attenuated ISR activation by the latter. ISRIB’s IC_50_ for attenuating Compound B activity, 23 nM (Fig. 6B), was in the range of ISRIB’s profile of action when inhibiting the ISR in stressed cells^12^. Attenuation of the Compound B-driven ISR was also observed at the level of mRNA translation, attesting to its “on pathway nature” (Fig. 6C and D). However, the plateau values of ISRIB’s effects on the ISR, ∼50% when antagonising Compound B and nearing 100% when antagonising stress-induced ISR activation^12^, remain a conspicuous and reproducible differential feature of the two responses to ISRIB.

**Figure 6:**
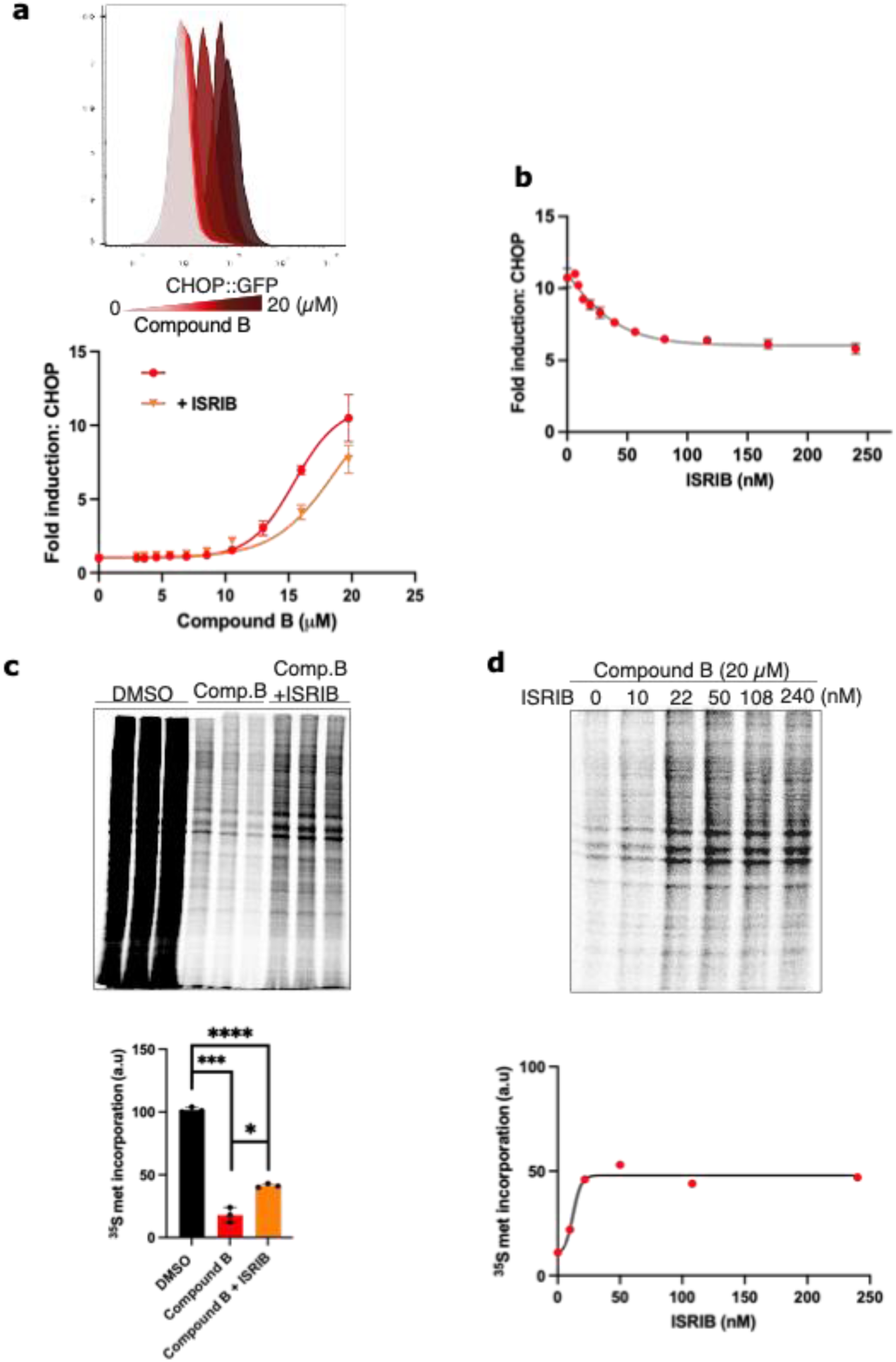
ISRIB partially attenuates Compound B action. **a,** Upper panel: Histogram of Compound B concentration-dependent CHOP::GFP activity measured by FACS. (shown is a representative experiment of 3). Lower panel: Fold increase (over untreated) of the median CHOP::GFP signal in pools of cells exposed for 18 hours to the indicated concentration of Compound B in the absence and presence of ISRIB (1 μM) (n = 3). **b,** Plot of the ISRIB-concentration dependent change in the CHOP::GFP signal of cells exposed to 20 μM Compound B (n = 3). The data were fitted by non-linear regression analysis to an inhibitor versus response model. IC_50_ = 22.83 nM (95% CI = 19.14-27.70). **c,** Upper panel: Autoradiograph of ^35^S methionine incorporation into newly synthesized cellular proteins in cells exposed to Compound B (20 μM) or Compound B + ISRIB (1 μM), performed in triplicate. Lower panel: quantification of the signal above. Statistical analysis was performed by a two-sided unpaired Welch’s t-test and significance is indicated by asterisks (* p<0.05) **d,** Upper panel: As in Fig. 5c, cells were exposed to Compound B (20 μM and the indicated concentration of ISRIB. Representative experiment of three. Lower panel: Quantification of ISRIB’s effect on ^35^S methionine incorporation into newly synthesized cellular proteins. The first and last data point, n= 3. The data were fitted by non-linear regression analysis to a dose response model using Prism GraphPad v10. (EC_50_ 12.58 nM; 95% CI = 0 – 27.94)

ISRIB action in cells is largely dependent on the presence of a pool of phosphorylated eIF2, whose inhibitory binding to eIF2B is antagonised allosterically by ISRIB engagement of eIF2B’s regulatory sites^10,11^. By contrast, Compound B retained substantial activity in *EIFS1* S51A mutant CHO cells, in which phosphorylation of serine 51 of eIF2α is precluded by its endogenous mutation to alanine^21^ (Extended Data Fig. 4A-C). Interestingly, ISRIB antagonism of Compound B action was lost in the eIF2αS51A mutant cells, suggesting that even when subjected to a direct eIF2B antagonist, ISRIB action requires the presence of a pool of phosphorylated eIF2.

In *EIF2B4 L180F* mutant cells with a structurally altered allosteric regulatory site due to a L180F mutation in the CHO eIF2Bδ subunit (the counterpart to human L179)^13,14^ Compound B activity was only modestly attenuated. This is reflected in the higher EC_50_ for CHOP::GFP activation [15 μM (14.94-16.21 95% CI) in wildtype CHO cells and 22.63 μM (20.88-26.69 μM, 95% CI) in *EIF2B4 L180F* mutant CHO cells] and in the extended tolerance of the mutant cells for the compound, which permitted its application at higher concentrations (Extended Data Fig. 4D-F). Reversal of the Compound B-induced ISR by ISRIB was lost in the eIF2BδL180F mutant cells, as expected of a mutation that interferes with ISRIB binding to eIF2B^20,26^. These features point to differences in target engagement by ISRIB and Compound B.

To assess the off-target effects of Compound B we monitored the activity of the IRE1/XBP1 axis, a stress response pathway that is activated in parallel to the ISR as endoplasmic reticulum stress mounts. The mRNA encoding the transcription factor XBP1 is rapidly spliced in a process initiated by the stress-activated ribonuclease, IRE1^22,23^. Exposure to thapsigargin rapidly depleted the pool of unspliced XBP1 and led to the accumulation of the spliced form (Extended Data Fig. 5). By contrast, Compound B minimally increased the level of spliced XBP1, without depleting the unspliced form. The modest increase in unspliced XBP1 caused by Compound B is consistent with an established cross-talk between the ISR and the IRE1/XBP1 axis, wherein the ISR increases total XBP1 mRNA^23^. The contrasting effects of thapsigargin (which activates both the IRE1 pathway and ISR) and Compound B point to the specificity of the latter towards the ISR.

Together these results show that Compound B, a symmetrical molecule based on Compound A-*(S)*, engages intracellular eIF2B and promotes robust ISR activation coherent with attenuation of translation initiation and a decline in protein synthesis modulated through availability of the ternary eIF2-GTP-Met-tRNA_i_ complex.

## Discussion

DEL screening with multiple conditions run in parallel is well suited to be designed to enable identification of compounds that engage their target with divergent MOAs. Its application here to eIF2B, led to the discovery of a novel compound that specifically interacted with eIF2B in complex with its inhibitory ligand, phosphorylated eIF2. Building on Compound A, the lead hit from the DEL screen, we synthesised an allosteric modulator of eIF2B, Compound B, that strongly activates the ISR. The potential existence of such a molecule was predicted based on first principles, but the discovery of Compound B, described here, is an experimental realisation of that prediction – the first that we are aware of.

The biophysical features of Compound A could not discriminate between direct stabilization of the eIF2B+eIF2(αP) complex by binding as a third component at the interface between eIF2B and the inhibitory phosphorylated α-subunit of eIF2 or binding to an allosteric site that stabilises the inhibitory engagement of eIF2(αP) indirectly by favouring the I-state of eIF2B. ISRIB probe displacement by Compound A was unhelpful in resolving the issue, given the antagonism between ISRIB binding and the I-state of eIF2B^10,11^. The issue was settled by the structure of the eIF2B+(P)eIF2α-NTD+Compound A-(*S*), which showed the latter to bind at a site far removed from the eIF2B+(P)eIF2α-NTD interface.

Due to the C2 symmetry of eIF2B when bound to two (P)eIF2 α-subunits, we were unable to determine orientation based on the position of Compound A-*(S)* in one half of the ISRIB-binding site. This explains the observation of density for the (trifluoromethyl)tetrahydronaphthalenyl)acetamide core of Compound A-*(S)* in both halves of the pocket and suggested an avenue for compound improvement through dimerisation.

Due to its poor solubility, we were unable to characterise the symmetric derivative of Compound A-(*S*), Compound B in vitro. Nonetheless features of the ancestral Compound A-(*S*) provide hints to its MOA. The binding site of Compound A-*(S)* was found to overlap that engaged by ISRIB. ISRIB and Compound A-*(S)* both lie in close proximity to eIF2BδL179 and eIF2BβN162 and by bridging two eIF2B βγδε tetramers to form an (βγδε)_2_ octamer both ISRIB^8^ and Compound A-(*S*) stabilise eIF2B in vitro. However, they diverge in other contacts, particularly with Compound A-*(S)*’s piperidine tail reaching into the core of eIF2B and interacting with both S227 residues of eIF2Bβ across the dimer interface. It is tempting to speculate that these divergent contacts account for the substantially different functional consequences of binding: stabilisation of the A-state by ISRIB that favours engagement of eIF2 as a substrate, thus enhancing GEF activity (and translation)^10,11^ and the opposite effect by Compound B, stabilisation of the I-state of eIF2B that favours engagement of eIF2(αP) as a negative regulator, thus attenuating GEF activity (and translation in vivo).

The effect of Compound A-(*S*) binding is mimicked by an H160D mutation in the beta subunit of eIF2B. This also biases the population of eIF2B towards the I-state. Notably, Compound A*-(S)* goes further than the partial I-state induction by H160D mutation and the I-state induced by Compound A-(*S*) was additionally supported by the salt bridging pattern as described by Lawrence et al.^18^ and comparison with previously published eIF2B + eIF2 complexes^9,14^.

Compound A-(*S*)’s MOA suggested the potential for ISR activation, realised by the synthesis of Compound B. The latter proved a powerful inducer of the ISR with plateau activation values similar to those attained by well-established activators such as thapsigargin and tunicamycin. This feature is mirrored by the plateau values of attenuated protein synthesis and the polysome profiles. However, Compound B’s potency is comparatively low, with EC_50_ values for ISR activation and protein synthesis inhibition of ∼10µM, likely a reflection of limited affinity for eIF2B.

When used at micromolar concentrations the potential for off-target effects of any compound is considerable. However, at concentrations associated with substantial inhibition of protein synthesis, Compound B had only modest effect on the parallel IRE1 mediated stress pathway. Therefore, whilst we cannot exclude engagement of targets other than eIF2B, we favour a parsimonious explanation whereby the binding of eIF2B at its allosteric site accounts for Compound B’s cellular effects and attribute the only partial reversal by ISRIB to different modes of binding by the two compounds that limit cross competition.

Protein synthesis is an essential function of cells and manipulations that strongly attenuate eIF2B GEF activity have fitness costs. This readily explains the toxicity observed with prolong application of high concentrations of Compound B to cells. Nonetheless when carefully regulated, partial inhibition of eIF2B’s GEF activity and the activation of the resulting downstream ISR is a powerful pro-survival feature. The compound described herein presents with a first-in-class eIF2B-directed ISR activation profile and the potential to magnify and extend the action of limited concentrations of eIF2(αP), meriting the appellation ISR activator, ISRAC. Compound B, as an ISRAC, represents an opposite of ISRIB, which preserves eIF2B GEF activity in the face of rising levels of eIF2(αP) and defends against the consequences of relentless ISR activation. Together these two tool compounds offer a promising foundation for the exploration of therapeutic strategies targeting diseases characterised by impaired ISR function. Activators of the ISR present as potential combination therapies in cancer to sensitize tumours and reduce growth^24,25^. Extending the ISR also has the potential to combat viral infection^26^ and certain protein misfolding diseases^27^. Further development of compounds with the MOA of Compounds A-(*S*) and B stands to benefit a comprehensive understanding of the molecular and translational consequences of ISR activation across diverse disease states and thus remains an important area for future investigation.

## Acknowledgements

We thank Reiner Schulte and Gabriela Grondys-Kotarba from the CIMR flow cytometry facility. This work was supported by AstraZeneca Ltd. D. R. is supported by the Wellcome Trust research grant 224407/Z/21Z. D. R. is a Wellcome Principal Research Fellow (224407/Z/21Z).

## Data availability

The cryo-EM map and model for the eIF2B + (P)eIF2α-NTD + Compound A-*(S)* structure were deposited to the EM Data Bank and PDB, respectively, under accession codes EMD-XXXXX, PDB-XXXX. Source data for Fig. 1, 2, 4 and 5 are provided with this paper.

## Contributions

1. T. M. O., D. R., D. B., P. B. and J. L. conceptualized the project and designed the experiments.

2. F. S., M. G-R. G. G., S. A., S. N. A., E. J. C. B., A. G., H. P. H., J. M. K., R. J. L., D. P., D. S., K. S., L. W., A. Z., carried out wet lab work or computational work for this project at AstraZeneca or CIMR. P. A. C., P. D., D. G., M-A. G., J. P. G., C. D. H., R. J., A. D. K., J. L., U. N., K. A. N., J. T. S. Y., Y. Y. and Y. Z. also carried out wet lab work or computational work for this project at X-Chem Inc.

3. T. M. O., D. R., D. B., J. L. and P. B. supervised the project and D. C., D. B., J. L. and P. B. established the collaboration with D. R.. E. R. oversaw the collaboration with X-Chem.

4. F. S. and M. G-R. wrote the initial draft and all authors contributed to the writing and revision of the paper and figures. D. R. contributed substantially to the editorial process of the paper.

## Corresponding authors

Correspondence to Fiona Shilliday or David Baker.

## Extended Data

- **Extended Data Fig. 1:** On-DNA ISRIB enrichment under expected conditions in the DEL screen
- **Extended Data Fig. 2:** Mass photometry QC of individual protein components and stabilization of the eIF2B complex upon addition of ISRIB and Compound A.
- **Extended Data Fig. 3:** Cryo-EM data processing
- **Extended Data Fig. 4:** Compound B retains its activity in *EIFS1Ser51Ala* cells and *EIF2B4Leu180Phe* cells
- **Extended Data Fig. 5:** Minimal activation of the IRE1/XBP1 axis in Compound B-treated cells

## Supplementary information

∉ **Supplementary Table 1: Cryo-EM data collection, refinement and validation statistics**
∉ **Supplementary Fig. 1:** Experimental infra-red spectra.
∉ **Supplementary Fig. 2:** Experimental VCD spectra.
∉ **Supplementary Fig. 3:** Overlay of the 41 lowest energy conformations (within 5 kJmol^-1^ of the minimum) used in the calculation of the Boltzmann average IR and VCD spectra.
∉ **Supplementary Table 2: Coordinates for the minimum energy conformation.**
∉ **Supplementary Fig. 4:** Comparison of calculated and experimental infra-red spectra.
∉ **Supplementary Fig. 5:** Comparison of calculated and experimental VCD spectra.

## Source data

- Source Data Fig. 1.
- Source Data Fig. 2.
- Source Data Fig. 4.
- Source Data Fig. 5.
- Source Data Fig. 6.

---

**Extended Data Fig. 1:**
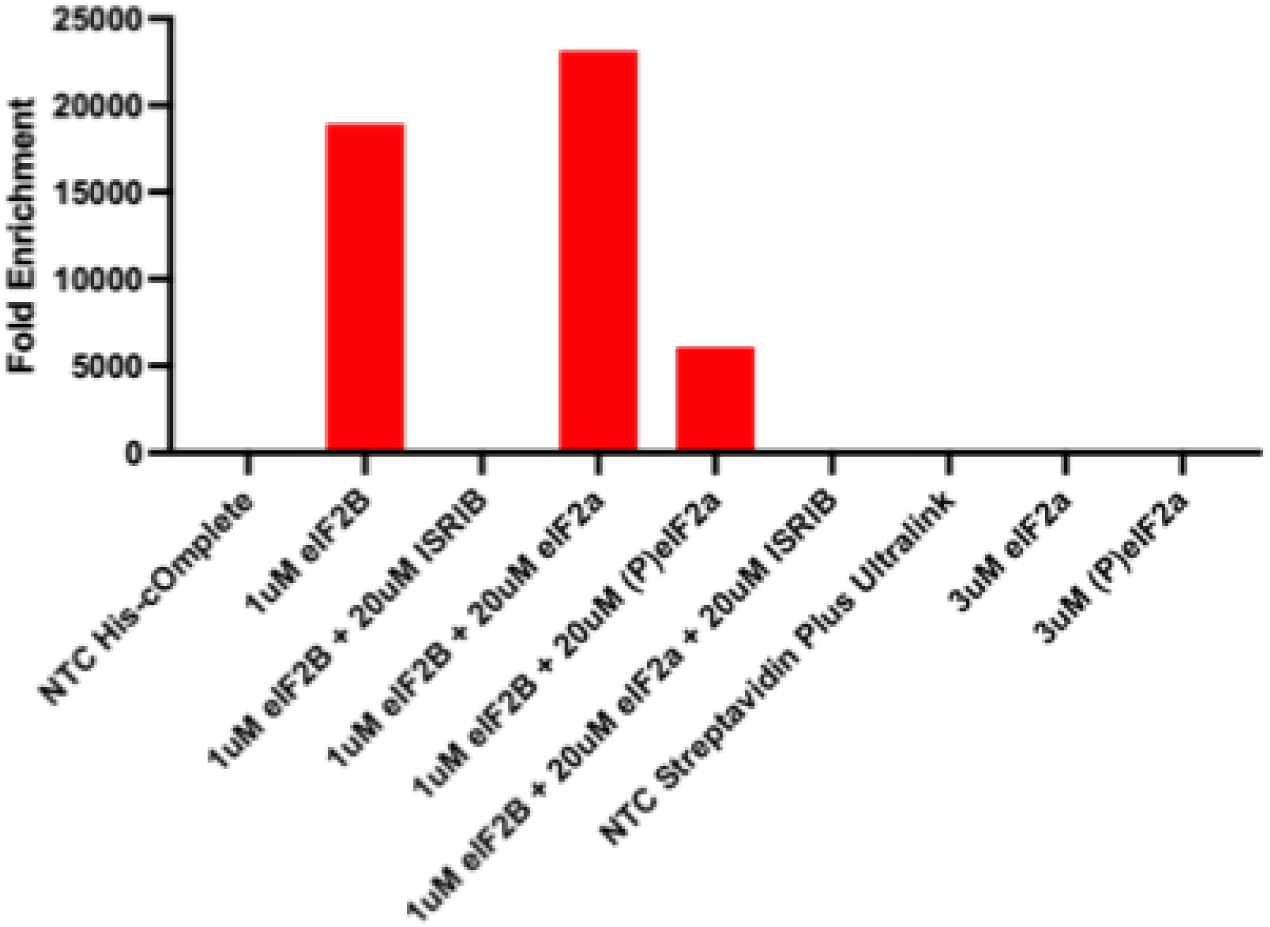
On-DNA ISRIB enrichment under expected conditions in the DEL screen. As a positive control on-DNA ISRIB was included in all conditions of the DEL screen. It showed significant enrichment under the expected conditions, binding to eIF2B alone, eIF2B + eIF2⍺ and to a lesser extent eIF2B + (P)eIF2⍺. Binding was reduced to non-significant levels in the parallel conditions when ISRIB was added to block the binding site. Enrichment was normalised to that of a corresponding DNA-tagged linker which was also included as an internal negative control in all conditions of the DEL screen.

**Extended Data Fig. 2:**
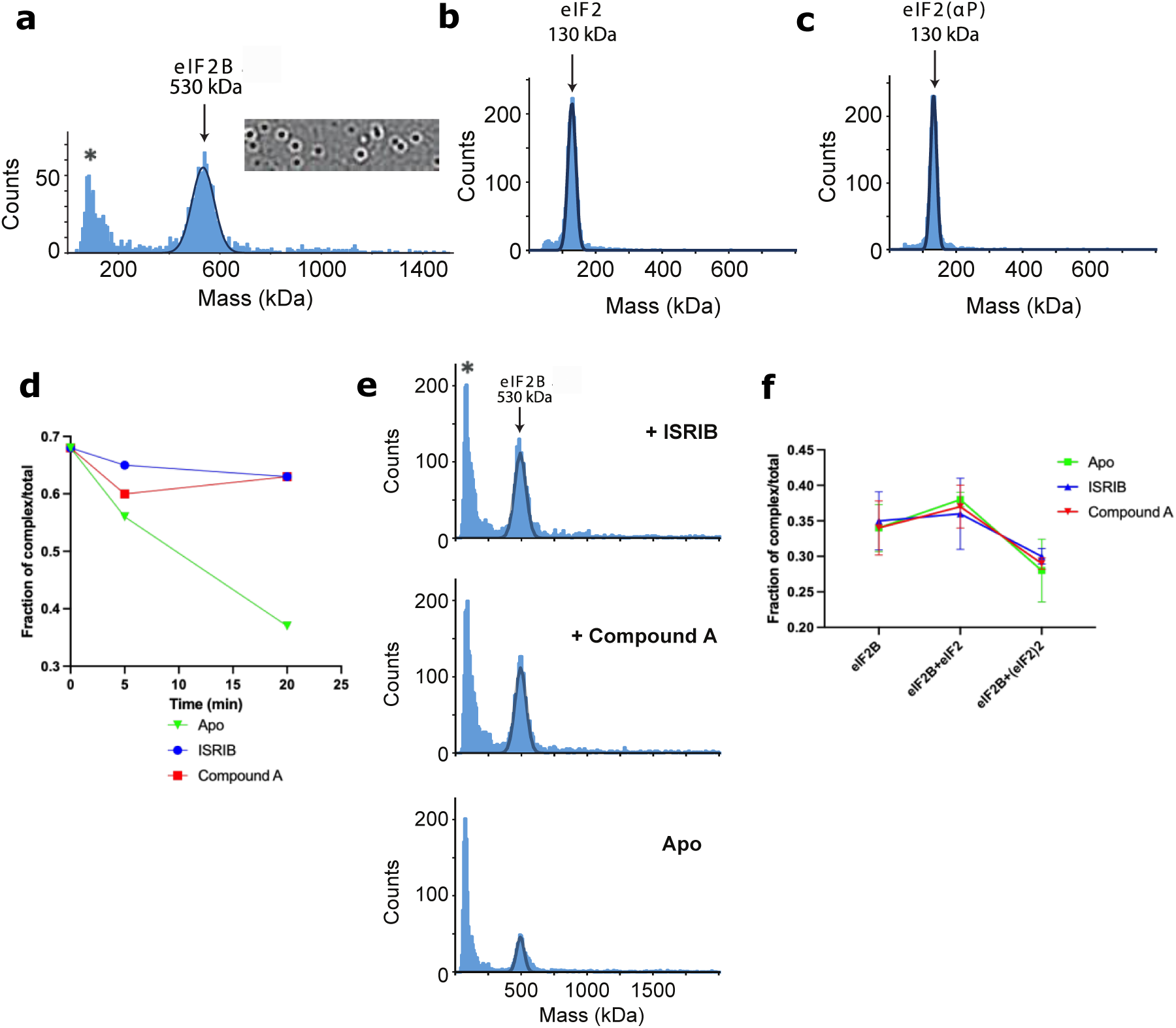
Mass photometry QC of individual protein components and stabilization of the eIF2B complex upon addition of ISRIB and Compound-A. **a, b, and c,** Mass photometry histograms of eIF2B (**a**), eIF2 (**b**) and eIF2(αP) (**c**) show samples are homogeneous and are of expected MWs (530, 130, 130 kDa, respectively). Note, the eIF2B sample also contains lower molecular weight species with a mass consistent with individual dissociated subunits and dimers thereof. (marked *) **d,** Time-dependent destabilization of eIF2B decamer in absence of ligand (Apo) or in presence of ISRIB or Compound A (33 μM) measured by mass photometry. **e,** Histograms of the distribution of the mass of the particles from the 20 min samples in ‘**d**’. As in **a** lower molecular weight species with a mass consistent with individual dissociated subunits and dimers thereof. (marked *) **f,** Mass photometry histograms of eIF2B-eIF2 complex in absence (Apo) or presence of either ISRIB or Compound A. Relative fraction of eIF2B, eIF2B-eIF2 and eIF2B-(eIF2)2 complexes in the absence or presence of compound.

**Extended Data Fig. 3:**
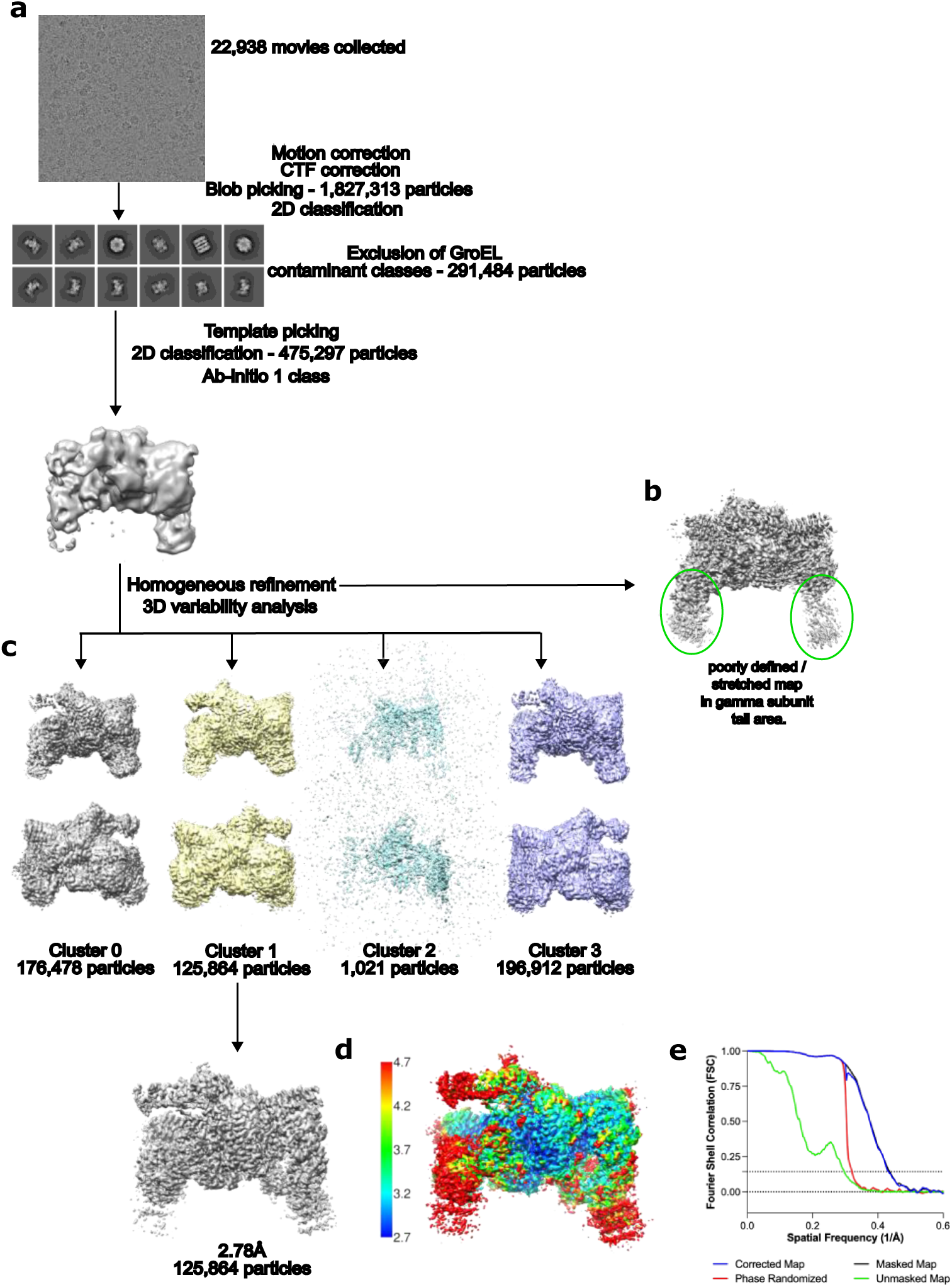
Cryo-EM data processing. **a,** Cryo-EM data processing strategy using cryoSPARC, movies collected and pre-processed in cryoSPARC Live, removing those with poor CTF fit or high amounts of motion, initial blob picking, then template picking to avoid picking GroEL particles seen clearly in the micrographs as a contaminant. 2D classification showing good quality classes of both eIF2B:(P)eIF2α-NTD complex and GroEL then ab-initio asking for 1 class. **b.** Intermediate homogeneous refinement map showing poorly defined / stretched map around the gamma subunit tails (green circles). **c,** 3D variability analysis was run following homogeneous refinement with 3 modes of variability. The 4 clusters requested are displayed with cluster 1 taken forward as it had improved density for 2 copies (P)eIF2α-NTD. Homogeneous refinement produced a consensus map of 2.67 Å resolution. **d,** Varying resolutions achieved across the cryo-EM map of eIF2B + (P)eIF2α + Compound A-*(S)* with high resolution (blue colours) in the core of eIF2B, but lower resolution in the outer subunits e.g. gamma and epsilon indicative of flexibility (red colours). **e,** Fourier Shell Correlation (FSC) for the final homogeneous refinement showing the 0.143 cut-off at a spatial frequency equivalent to 2.67 Å.

**Extended Data Figure 4:**
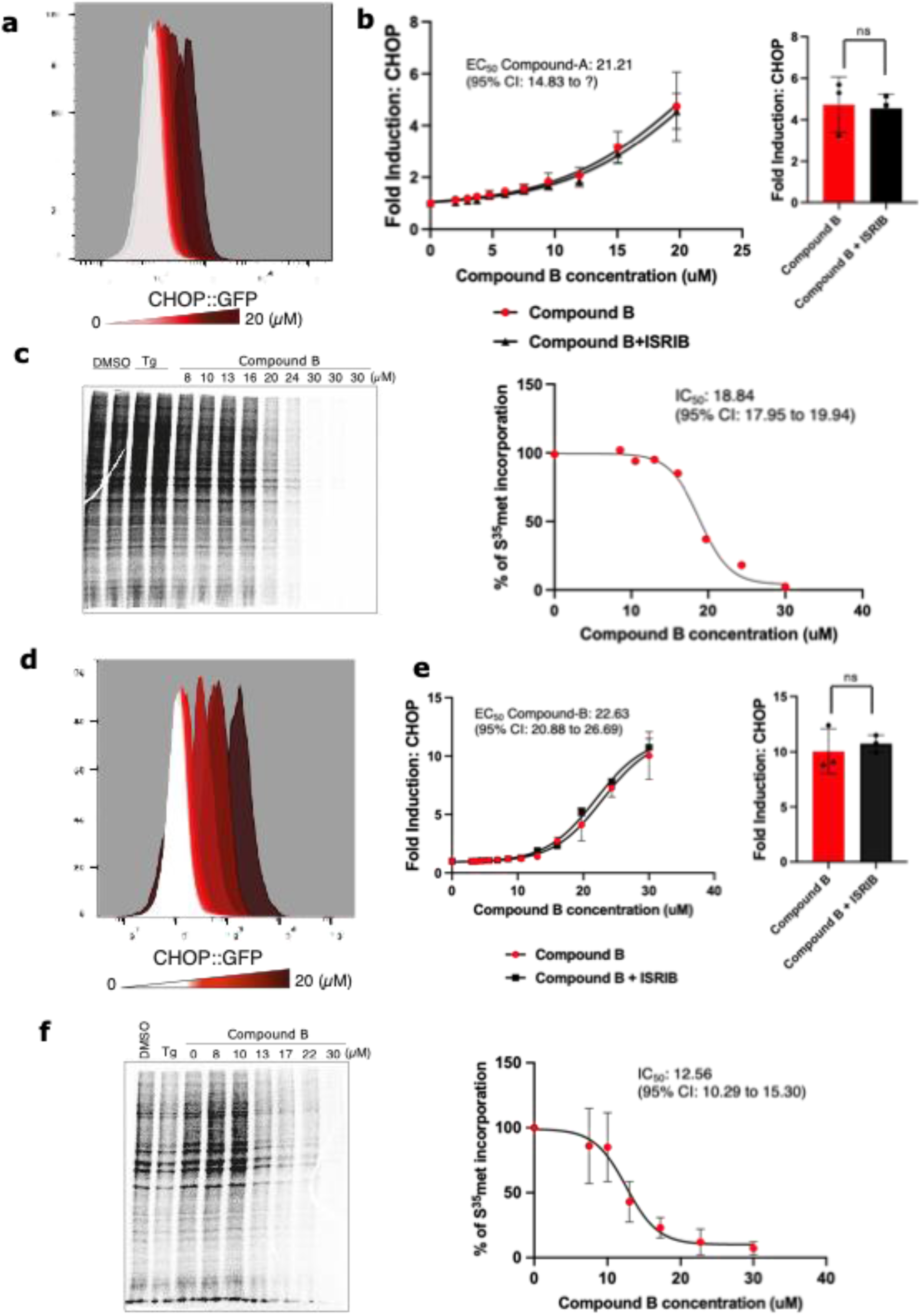
Compound B retains its activity in *EIFS1^Ser51Ala^* cells and *EIF2B4^Leu^*^180^*^Phe^* cells. **a,** Histogram of CHOP::GFP ISR reporter gene activity (measured by FACS) in *EIFS1^Ser51Ala^* cells exposed for 18 hours to the indicated concentration of Compound B. **b,** Median CHOP::GFP activity derived from the histogram as fold increase over the level in untreated cells, as a function of Compound B concentration. Shown is n = 3. The data were fitted by non-linear regression analysis to a dose response model. EC50 values with 95% confidence intervals are shown. Also shown is reporter activity in the presence of 1 μM ISRIB. The bar chart to the right shows the mean ± SD of the fold increase (over untreated) of samples exposed to compound B (20 μM) in presence and absence of ISRIB (1 μM). Statistical analysis was performed by a two-sided unpaired Welch’s t-test (ns: non-significant). **c,** Left panel: Autoradiograph of an SDS-PAGE of lysates from *EIFS1^Ser51Ala^* cells incorporating ^35^S methionine over a 30 min period following exposure to the indicated concentration of Compound B commencing 30 minutes before labelling and continued throughout the labelling period. Shown is a representative from three such experiments performed. Right panel: ^35^S methionine incorporation plotted as a function of Compound B concentration. The data were fitted by non-linear regression analysis to an inhibitor versus response model. IC_50_ values with 95% confidence intervals are shown. **d, & e,** As in **a,** and **b,** but for CHOP::GFP ISR reporter gene in *EIF2B4^Leu^*^180^*^Phe^* cells. **f,** Left panel: Autoradiograph of an SDS-PAGE of lysates from *EIF2B4^Leu^*^180^*^Phe^* cells incorporating ^35^S methionine over a 30 min period following exposure to thapsigargin (1 μM) or the indicated concentration of Compound B for 30 min. Shown is a representative from three such experiments performed. Right panel: ^35^S methionine incorporation plotted as a function of Compound B concentration. The data were fitted by non-linear regression analysis to a dose response model. IC_50_ values with 95% confidence intervals are shown.

**Extended Data Figure 5:**
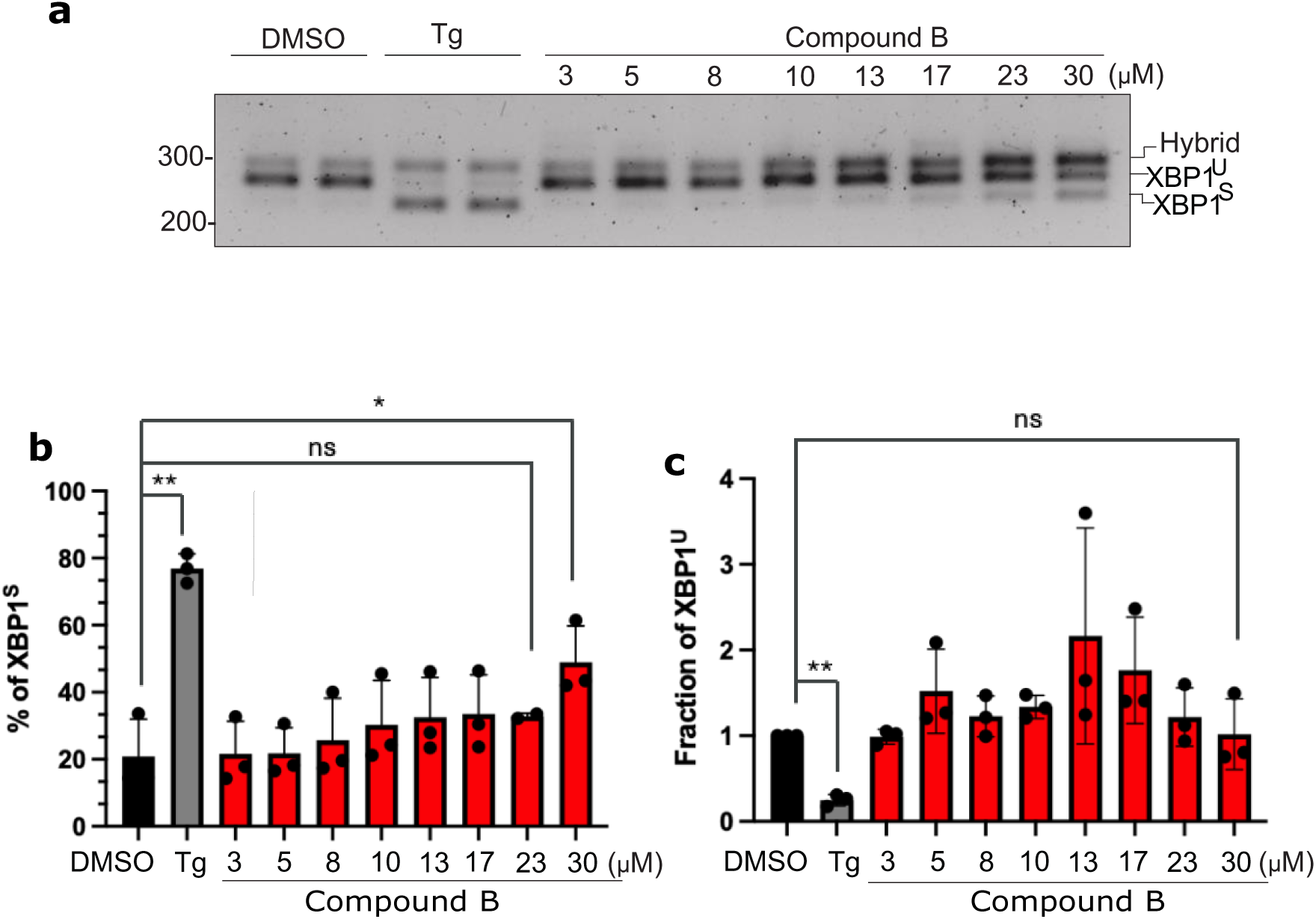
Minimal activation of the IRE1/XBP1 axis in Compound B-treated cells. **a,** Photomicrograph of a stained agarose gel resolving isoforms of XBP1 cDNA derived by reverse-transcriptase PCR (RT-PCR) from lysates of cells exposed for 1h to Tg (1 μM) or the indicated concentration of Compound B. The migration of the different isoforms is indicated: XBP1^S^ and XBP1^U^ are duplexes of the spliced and unspliced XBP1 cDNA and the hybrid species of both, generated in the RT-PCR procedure is noted. Shown is a gel representative of three such experiments performed. **b, and c,** quantitation of the relative abundance of the spliced and unspliced cDNA in each sample. Shown are the means and SD well as individual data points of the three replicates. Statistical analysis was performed by a two-sided unpaired Welch’s t-test and significance is indicated by asterisks (*p <0.05; ** p < 0.01; ns: non-significant).

## Extra references for methods

Adomavicius et al., 2019^28^

Punjani, et al. 2017^29^

Punjani and Fleet 2021^30^

Emsley et al. 2010^31^

Grade, version 1.2.20^32^

Liebschner et al. 2019^33^

Pettersen et al. 2004^34^

PyMOL Molecular Graphics System, Version 2.0 Schrödinger, LLC.^35^

Novoa et al. 2001^36^

Sekine et al. 2016^37^

Ordóñez et al., 2021^38^

## References

1. Pakos-Zebrucka, K., et al. The integrated stress response. EMBO Rep. 17, 1374–1395 (2016).

2. Donnelly, N., Gorman, A. M., Gupta, S. & Samali, A. The eIF2α kinases: their structures and functions. Cell. Mol. Life Sci. 70, 3493–3511 (2013).

3. Hinnebusch, A. G., Ivanov, I. P. & Sonenberg, N. Translational control by 5′-untranslated regions of eukaryotic mRNAs. Science 352, 1413–1416 (2016).

4. Harding, H. P. et al. Regulated translation initiation controls stress-induced gene expression in mammalian cells. Mol. Cell 6, 1099–1108 (2000).

5. Vattem, K. M. & Wek, R. C. Reinitiation involving upstream ORFs regulates ATF4 mRNA translation in mammalian cells. Proc. Natl. Acad. Sci. U. S. A. 101, 11269–11274 (2004).

6. Costa-Mattioli, M. & Walter, P. The integrated stress response: From mechanism to disease. Science 368, eaat5314 (2020).

7. Hanson, F. M., Hodgson, R. E., de Oliveira, M. I. R., Allen, K. E. & Campbell, S. G. Regulation and function of elF2B in neurological and metabolic disorders. Biosci. Rep. 42, BSR20211699 (2022).

8. Tsai, J. C. et al. Structure of the nucleotide exchange factor eIF2B reveals mechanism of memory-enhancing molecule. Science 359, eaaq0939 (2018).

9. Kashiwagi, K. et al. Structural basis for eIF2B inhibition in integrated stress response. Science 364, 495– 499 (2019).

10. Zyryanova, A. F. et al. ISRIB Blunts the Integrated Stress Response by Allosterically Antagonising the Inhibitory Effect of Phosphorylated eIF2 on eIF2B. Mol. Cell 81, 88–103.e6 (2021).

11. Schoof, M. et al. eIF2B conformation and assembly state regulate the integrated stress response. eLife 10, e65703 (2021).

12. Sidrauski, C. et al. Pharmacological brake-release of mRNA translation enhances cognitive memory. eLife 2, e00498 (2013).

13. Sekine, Y. et al. Mutations in a translation initiation factor identify the target of a memory-enhancing compound. Science 348, 1027–1030 (2015).

14. Zyryanova, A. F. et al. Binding of ISRIB reveals a regulatory site in the nucleotide exchange factor eIF2B. Science 359, 1533–1536 (2018).

15. Wong, Y. L. et al. eIF2B activator prevents neurological defects caused by a chronic integrated stress response. eLife 8, e42940 (2019).

16. Craig, R. A. et al. Discovery of DNL343: A Potent, Selective, and Brain-Penetrant eIF2B Activator Designed for the Treatment of Neurodegenerative Diseases. J. Med. Chem. 67, 5758–5782 (2024).

17. Boone, M. et al. A point mutation in the nucleotide exchange factor eIF2B constitutively activates the integrated stress response by allosteric modulation. eLife 11, e76171 (2022).

18. Lawrence, R. E. et al. A helical fulcrum in eIF2B coordinates allosteric regulation of stress signaling. Nat. Chem. Biol. 20, 422–431 (2024).

19. Harding, H. P. et al. Bioactive small molecules reveal antagonism between the integrated stress response and sterol-regulated gene expression. Cell Metab. 2, 361–371 (2005).

20. Harding, H. P., Zhang, Y. & Ron, D. Protein translation and folding are coupled by an endoplasmic-reticulum-resident kinase. Nature 397, 271–274 (1999).

21. Crespillo-Casado, A., Chambers, J. E., Fischer, P. M., Marciniak, S. J. & Ron, D. PPP1R15A-mediated dephosphorylation of eIF2α is unaffected by Sephin1 or Guanabenz. eLife 6, e26109 (2017).

22. Yoshida, H., Matsui, T., Yamamoto, A., Okada, T. & Mori, K. XBP1 mRNA is induced by ATF6 and spliced by IRE1 in response to ER stress to produce a highly active transcription factor. Cell 107, 881–891 (2001).

23. Calfon, M. et al. IRE1 couples endoplasmic reticulum load to secretory capacity by processing the XBP-1 mRNA. Nature 415, 92–96 (2002).

24. Mijit, M. et al. Activation of the integrated stress response (ISR) pathways in response to Ref-1 inhibition in human pancreatic cancer and its tumor microenvironment. Front. Med. 10, (2023).

25. Cyran, A. M. et al. Inhibition of EIF2α Dephosphorylation Decreases Cell Viability and Synergizes with Standard-of-Care Chemotherapeutics in Head and Neck Squamous Cell Carcinoma. Cancers 15, 5350 (2023).

26. Wuerth, J. D. & Weber, F. Shielding the mRNA-translation factor eIF2B from inhibitory p-eIF2 as a viral strategy to evade protein kinase R-mediated innate immunity. Curr. Opin. Immunol. 78, 102251 (2022).

27. D’Antonio, M. et al. Resetting translational homeostasis restores myelination in Charcot-Marie-Tooth disease type 1B mice. J. Exp. Med. 210, 821–838 (2013).

28. Adomavicius, T. et al. The structural basis of translational control by eIF2 phosphorylation. Nat. Commun. 10, 1–10 (2019).

29. Punjani, A., Rubinstein, J. L., Fleet, D. J. & Brubaker, M. A. cryoSPARC: algorithms for rapid unsupervised cryo-EM structure determination. Nat. Methods 14, 290–296 (2017).

30. Punjani, A. & Fleet, D. J. 3D variability analysis: Resolving continuous flexibility and discrete heterogeneity from single particle cryo-EM. J. Struct. Biol. 213, 107702 (2021).

31. Emsley, P., Lohkamp, B., Scott, W. G. & Cowtan, K. Features and development of Coot. Acta Crystallogr. D Biol. Crystallogr. 66, 486–501 (2010).

32. Smart, O. S. et al. Grade, version 1.2.20. Cambridge, United Kingdom, Global Phasing Ltd., https://www.globalphasing.com. 2011,.

33. Liebschner, D. et al. Macromolecular structure determination using X-rays, neutrons and electrons: recent developments in Phenix. Acta Crystallogr. Sect. Struct. Biol. 75, 861–877 (2019).

34. Pettersen, E. F. et al. UCSF Chimera--a visualization system for exploratory research and analysis. J. Comput. Chem. 25, 1605–1612 (2004).

35. The PyMOL Molecular Graphics System, Version 3.0 Schrödinger, LLC.

36. Novoa, I., Zeng, H., Harding, H. P. & Ron, D. Feedback inhibition of the unfolded protein response by GADD34-mediated dephosphorylation of eIF2alpha. J. Cell Biol. 153, 1011–1022 (2001).

37. Sekine, Y. et al. Paradoxical Sensitivity to an Integrated Stress Response Blocking Mutation in Vanishing White Matter Cells. PloS One 11, e0166278 (2016).

38. Ordóñez, A., Harding, H. P., Marciniak, S. J. & Ron, D. Cargo receptor-assisted endoplasmic reticulum export of pathogenic α1-antitrypsin polymers. Cell Rep. 35, 109144 (2021).

