## Supplemental methods for "A molecular stabiliser of an inhibitory eIF2B-eIF2(αP) complex activates the Integrated Stress Response"

### Materials and methods

#### Protein expression and purification of recombinant proteins

Molecular cloning, expression and purification of human eIF2B

The molecular cloning strategy followed for the expression of human eIF2B was based on previously reported studies[8, 11]. In summary, the DNA sequence coding for human *eIF2B1* (UniProt accession: Q14232) was cloned into pET28c+ expression vector (GenScript) to produce N-terminal 6His-3C-tagged eIF2Bα protein. For DNA-encoded library screening, the N-terminal modification was replaced with a 6His-tobacco etch virus (TEV) protease site (ENLYFQ/GS) tag. *eIF2B*2 (P49770) and *eIF2B4* (Q9UI10) were inserted into sites 1 and 2 of pETDuet-1 (GenScript), respectively. *eIF2B3* (Q9NR50) and *eIF2B5* (Q13144) were cloned into sites 1 and 2 of pCOLADuet-1 (GenScript), respectively, to produce N-terminal 6His-3C-tagged eIF2B∑ protein. For HTRF studies, an AVI tag was inserted to the existing eIF2B∑ modification to enable site-specific biotinylation. All the previous expression plasmids were codon-optimised for expression in *E. coli* and purchased from Genscript. The appropriate DNA constructs necessary for eIF2B expression and decamer self-assembly were cotransformed into chemically competent *E. coli* BL21 (DE3) Gold cells (New England Biolabs) and grown at 37 °C with shaking. Protein overexpression was induced by addition of 0.1 mM IPTG at log phase cultures (0.6-0.8 $OD_{600}$) and incubated for a further 20 h at 18 °C. Cells were harvested by centrifugation and stored at –80 °C. The cell pellet was resuspended in Lysis Buffer (20 mM HEPES, pH 7.5, 250 mM KCl, 15 mM Imidazole, 5 mM $MgCl_2$, 1 mM TCEP, cOmplete EDTA-free Protease Inhibitor Tablets (Roche) and benzonase nuclease) and passed twice through a Constant Systems cell homogenizer at 25-27 kpsi. The lysate was clarified by 1 h 20 min centrifugation at 16,000 rpm and incubated overnight with TALON metal resin (Takara) pre-equilibrated in Wash Buffer (20 mM HEPES, pH 7.5, 200 mM KCl, 15 mM Imidazole, 5 mM $MgCl_2$ 1 mM TCEP) for $Co^{2+}$-affinity purification in batch mode. The protein was eluted off the column using Wash Buffer containing 200 mM Imidazole. Next, the eIF2B fraction was diluted 1:10 in AIEX Buffer (20 mM HEPES, pH 7.5, 5 mM $MgCl_2$ 1 mM TCEP) and immediately loaded into a 5 mL HiTrap Q HP column washed in the same buffer using an ÄKTA Pure (Cytiva) system at 4 °C. The eIF2B fractions eluted off the column at 30 mS/cm conductivity using a linear gradient to 700 mM KCl. AVI tagged eIF2B complex for HTRF was subjected to biotinylation using BirA biotin-protein ligase as per manufacturer's protocol (Avidity). eIF2B complex for DEL was processed with 300 U of HRV 3C-protease (Sigma-Aldrich) and incubated overnight at 4 °C for tag cleavage. Finally, the eIF2B sample was concentrated and applied to a HiLoad 26/600 Superdex 200 pg size-exclusion column (Cytiva) equilibrated in 20 mM HEPES, pH 7.5, 200mM KCl, 5 mM $MgCl_2$ 1 mM TCEP. Peak fractions were concentrated and aliquots flash-frozen in liquid nitrogen. Protein identity and labelling were confirmed by intact mass spectrometry using a Sciex X500B Q-TOF with Sciex Excion LC instrument and bioZen 3.6 µm Intact XB-C8 column. All proteins were diluted at least 10X in mass spectrometry buffer (5 % acetonitrile, 0.1 % formic acid) to 0.1 mg/ml.

<u>Molecular cloning, expression and purification of human eIF2 heterotrimer and eIF2〈-NTD</u>

The generation of eIF2 heterotrimer was adapted from previously reported studies[9]. DNA fragments coding for *EIF2S1* (P05198), C-terminal TEV cleavage site-3xFLAG-8His tagged *EIF2S2* (P20042) and *EIF2S3* (P41091) were synthesized by GeneArt and cloned individually into pMAZ vectors via Golden Gate assembly for cotransfection into Expi293F cells. Cells were harvested after 48h transient protein expression and resuspended in 20 mM HEPES, pH 6.8, 150 mM KCl, 20 mM Imidazole, 1 mM $MgCl_2$, 0.5 mM EDTA, 10 %(v/v) glycerol, 0.1 %(v/v) Triton X-100, 5 mM β-mercaptoethanol (BME) and cOmplete EDTA-free Protease Inhibitor Tablets (Roche) for lysis using a Constant Systems cell homogenizer at 15 kpsi. The lysate was clarified by 1 h 30 min centrifugation at 16,000 rpm and applied to a HisTrap HP 5 ml $Ni^{2+}$ column (Cytiva) for affinity purification. The column was washed with Wash Buffer (20 mM HEPES, pH 6.8, 150 mM KCl, 20 mM Imidazole, 1 mM $MgCl_2$, 10 %(v/v) glycerol, 5 mM BME) and $Ni^{2+}$-binding protein was eluted off the column using Wash Buffer containing 500 mM Imidazole. The same procedure was repeated using the flow-through to maximise protein recovery. Elution fractions were pooled, dialysed extensively against 4 L Wash Buffer and processed for eIF2® C-terminal tag removal using 6His-TEV protease. The dialysed sample was passed back over $Ni^{2+}$ for subtractive affinity. The cleaved protein was diluted 1:3 with AIEX A Buffer (20 mM HEPES, pH 6.8, 100 mM KCl, 1 mM $MgCl_2$, 10 %(v/v) glycerol, 5 mM BME) and purified by ion-exchange chromatography using a HiTrap Q 5 ml column (Cytiva). The eIF2 fractions eluted off the column at 20 mS/cm conductivity using a linear gradient to 1 M KCl. When appropriate, the eIF2 sample was processed for *in vitro* phosphorylation using recombinant kinase domain of PERK protein. Finally, the sample was applied to a HiLoad 16/600 Superdex 200 pg column (Cytiva) equilibrated in 20 mM HEPES, pH 7.5, 100 mM KCl, 0.1 mM $MgCl_2$, 10 %(v/v) glycerol, 1 mM TCEP. The fractions containing eIF2 heterotrimer were concentrated, aliquoted and stored at –80 °C until use. Protein identity was confirmed by LC/MS.

The generation of C-terminal AVI tagged N-terminal domain of eIF2α$^{1-187}$ (eIF2α-NTD) was based on previous studies[10]. In summary, human eIF2〈$^{1-187}$ containing a N-terminal 6His-TEV sequence and C-terminal AVI tag was cloned into azET01 vector using NdeI/XhoI restriction sites (NBS Biologicals) and optimised for *E. coli* expression. Following transformation into BL21 (DE3) Gold *E. coli* cells, protein overexpression was induced at log phase cultures and induced with 0.1 mM IPTG. Pelleted cells were resuspended in 20 mM HEPES, pH 7.5, 250 mM NaCl, 40 mM Imidazole, 1 mM TCEP, cOmplete EDTA-free Protease Inhibitor Tablets (Roche) and benzonase nuclease and lysed using by high pressure pulses at 25 kpsi. The clarified lysate was purified by Ni-NTA affinity purification in batch mode and the eluate subjected to 6His-TEV protease cleavage. The sample was dialysed to remove high levels of imidazole, purified by subtractive affinity and processed with BirA biotin-protein ligase (Avidity) for site-specific biotinylation. Finally, the protein was loaded onto a HiLoad 16/600 Superdex 75pg column (Cytiva) stored in 50 mM HEPES, pH 7.5, 200 mM KCl, 1 mM TCEP for gel filtration, aliquoted and stored at –80 °C. Protein identity and labelling were confirmed by LC/MS.

<u>eIF2 fluorescent nucleotide loading and eIF2〈 S51 phosphorylation</u>

Loading of GDP-BODIPY to eIF2 for guanine nucleotide exchange assays was performed according to Wong et al., 2018[15].

eIF2〈 S51 phosphorylation of eIF2 heterotrimer and eIF2α-NTD was performed either *in vitro* using purified recombinant kinase domain of PERK$^{588-1116}$ or *in vivo* (eIF2α-NTD only) by cotransformation and expression in *E. coli* BL21 (DE3) Gold cells. To enable both approaches, DNA fragments coding for N-terminal GST tagged *EIF2AK3* (<u>Q9NZJ5</u>) corresponding to amino acid residues R588-N1116 were synthesized by GeneArt and cloned into a pACYC vector via Golden Gate assembly. For recombinant PERK kinase domain protein production, the expression vector was transformed into *E. coli* BL21 (DE3) Gold cells and grown at 37 °C with shaking. Protein overexpression was induced by addition of 0.5 mM IPTG at log phase cultures (0.6-0.8 $OD_{600}$) and incubated for a further 20 h at 18 °C. Cells were harvested by centrifugation and stored at -80°C. The cell pellet was resuspended in 50 mM HEPES, pH 7.5, 300 mM NaCl, 1 mM TCEP, cOmplete EDTA-free Protease Inhibitor Tablets (Roche) and benzonase nuclease followed by lysis using a Constant Systems cell homogenizer at 25-27 kpsi. After centrifugation, the clarified lysate was loaded onto a GSTrap HP 5ml column (Cytiva) for affinity purification. The column was washed with 50 mM HEPES, pH 7.5, 300 mM NaCl, 1 mM TCEP and GST-tagged protein eluted with 10 mM reduced glutathione (Sigma Aldrich). The eluate was subjected to size-exclusion chromatography using a HiLoad 16/600 Superdex 75 pg size-exclusion column (Cytiva) equilibrated in the same buffer with no reduced glutathione. The pooled fractions containing $PERK^{588\text{-}1116}$ were concentrated, aliquoted and stored at –80 °C until use.

For *in vitro* phosphorylation, the eIF2 reagents were mixed with $PERK^{588\text{-}1116}$ at a 500:1 molar ratio, respectively, in Phospho Buffer (100 mM HEPES pH 7.4, 750 mM KCl, 5 mM TCEP, 10 mM $MgCl_2$, 2 mM ATP) and incubated at room temperature for 2h. Phosphorylated eIF2 reagents were confirmed by intact LC/MS and Western Blot.

<u>Western blotting</u>

Protein reagents were run on 4-12 % SDS-PAGE gel, transferred onto a nitrocellulose membrane and blocked using PBS Blocking Buffer (LI-COR Biosciences) incubating for 1h at room temperature. Next, the membrane was incubated with primary monoclonal rabbit anti-(P)eIF2α(Ser51) antibody (#9721, Cell Signaling Technology) at 4 °C overnight. Subsequent steps were adapted from Adomavicius et al., 2019[28]. Signal was then captured by using Bio-Rad ChemiDoc imager.

<u>Complex formation of eIF2B:(P)eIF2α-NTD bound to Compound A-*(S)* for Cryo-EM</u>

For complex assembly, (P)eIF2α-NTD was added to purified eIF2B in a 4-fold molar excess in a final solution containing 20 mM HEPES, pH 7.5, 100 mM KCl, 1 mM $MgCl_2$, 1 mM TCEP and incubated for 1h at room temperature. For eIF2B:(P)eIF2α-NTD formation bound to the Compound A-*(S)*, a 120-fold molar excess of compound compared to eIF2B was added to the reaction mix, and a 100 μM final concentration was kept across the storage buffer. After incubation, protein complexes were purified using a Superose 6 Increase 10/300 GL column (Cytiva), and fractions corresponding to eIF2B:(P)eIF2α-NTD bound to Compound A-*(S)* were pooled, concentrated and flash-frozen in liquid nitrogen.

**<u>DEL screening</u>**

Twenty-six unique DNA-encoded libraries (DELs) were mixed together and incubated with 1 μM His-eIF2B in 60 μL of Selection Buffer (50 mM Tris pH 7.5, 150 mM NaCl, 2 mM $MgCl_2$, 1 mM DTT, 0.01 % Triton X-100, 1 mg/mL sssDNA). Separate samples were prepared to enable the parallel screening of this library deck under multiple conditions, including a no-target control, 1 μM His-eIF2B, 1 μM His-eIF2B pre-incubated with 20 μM ISRIB, 1 μM His-eIF2B in complex with eIF2α, and 1 μM His-eIF2B in complex with (P)eIF2α. Biotinylated eIF2α and Biotinylated (P)eIF2α were prepared as counter-screens to informatically filter out hits which do not bind directly to eIF2B. Binding partners were pre-incubated with eIF2B for 30 minutes prior to addition of 1 nanomole of the DNA-encoded library mix, 3 attomoles of the positive control DNA-tagged ISRIB, and 30 attomoles of a corresponding negative control DNA lacking any conjugated small molecule. After a one-hour incubation of this mixture, the proteins and associated library members were captured onto 5 μL of pre-equilibrated His-cOmplete Phytip using the Phynexus ME200 automated multi-channel pipettor. Proteins were incubated with the matrix for 30 minutes, matrices were washed 8 times with selection buffer, and bound library members were eluted by heating matrices to 85 °C for 5 minutes in selection buffer. A second round of selection was performed to further enrich the hits using fresh proteins and other reagents and the round one output. The screening output from round two was then PCR amplified using primers which contain READ1 and READ2 sequences for Illumina sequencing. Sequencing was performed on an Illumina 2500 instrument in high-output mode, yielding 722 million single-end reads across all selection conditions and libraries for an average of 3.1 million reads per library per condition.

**Mass photometry**

Mass photometry was performed on a Refeyn One instrument (Refeyn Ltd, UK). All samples were diluted in HBS-N (10 mM HEPES pH 7.4, 150 mM NaCl) and analyzed using 60 s acquisition time. Peak contrast was calculated from the resulting histograms using Gaussian fits. The contrast-to-mass conversion was achieved using a native protein ladder with known species of different sizes (NativeMark, ThermoFisher). Protein complexes were measured individually to confirm sample homogeneity and size. eIF2B complex $(\alpha\beta\gamma\delta\varepsilon)_2$, eIF2 and eIF2(αP) were rapidly diluted from stock concentration (4.5 μM, 41 μM or 61 μM respectively) to ~50 nM to enable accurate measurements. The same measurement principle was applied for the time-resolved measurement with the addition of compound (33 μM). Similarly, eIF2B in complex with eIF2 or eIF2(αP) was incubated at 1:2 ratio at 1 μM eIF2B before rapid dilution to 50 nM upon measurement in absence or presence of compound (33 μM).

**Biolayer interferometry (BLI) assay**

BLI experiments were conducted on an Octet RED instrument (ForteBio) set at 25°C and an orbital shake speed of 1000rpm, using Streptavidin-coated biosensors (ForteBio/Sartorius catalog # 18-5019) and 96-well plates (Greiner F bottom #655209) with a volume per well of 200 μl. The method was setup using the Octet Data Acquisition software version 6.4. The assay buffer was composed of 20 mM HEPES pH 7.5, 150 mM KCl, 2 mM $MgCl_2$, 0.01% Triton X-100, 1 mM TCEP, 0.05 mg/ml BSA and 1% DMSO.

Biosensors were pre-incubated for a minimum of 10 minutes in assay buffer prior to the start of the experiment. Each compound was tested in the dissociation step of an assay defined as follows. Biosensors were pre-conditioned by dipping them into a 10 mM Glycine pH 1.7 solution for 5s followed by 5s in assay buffer repeated 5 times. They were then equilibrated in assay buffer for 60s prior to loading of 121 nM biotinylated (P)eIF2α-NTD to a binding signal of 0.8 nm. Free streptavidin sites were blocked for 120s using a 50 μM EZ-Link™ Amine-PEG2-Biotin solution, followed by a 60s baseline equilibration step in assay buffer. Association of eIF2B diluted to 66.5nM was monitored for 500s, and dissociation of eIF2B was measured for 1000s in assay buffer containing either DMSO or a 7-point serial dilution of compound (0.39 – 25 μM, 1:2 dilution step) prepared using an HP D300 dispenser.

Raw data were exported as .csv files from the ForteBio Data Analysis software version 6.4, processed using Microsoft Excel version 2102 and analysed using GraphPad Prism 8 (version 8.4.3).

The dissociation traces were aligned to the end of the baseline step preceding the association step and normalized to the binding signal at the end of the association step, corresponding to the amount of biotinylated (P)eIF2α-NTD:eIF2B formed.

The normalized dissociation traces were fitted to a biphasic exponential decay model, using the equations:

SpanFast = (Y0-Plateau)*PercentFast*.01

SpanSlow = (Y0-Plateau)*(100-PercentFast)*.01

Y = Plateau+SpanFast*exp(-KFast*X)+SpanSlow*exp(-KSlow*X)

where Y0 is the Y value when X (time) is zero, Plateau is the Y value at infinite times, Kfast and Kslow are the two rate constants expressed in reciprocal of the X axis time units, PercentFast is the fraction of the span (from Y0 to Plateau) accounted for by the faster of the two components.

The following constraints were applied to the fit: KFast and KSlow were both shared between all datasets, and the Plateau was set to a constant value of 0.

To determine the compound $EC_{50}$, the PercentFast values extracted from the fits of the dissociation traces were normalized to the PercentFast value of the dissociation recorded in absence of compound. The normalized PercentFast values were plotted against the concentration of compound and the data were fitted using the [Inhibitor] vs. Response (three parameters, nHill=-1) model, to the following equation:
Y=Bottom + (Top-Bottom)/(1+(X/EC50))
where EC50 is the concentration of compound that gives a response halfway between Bottom and Top, Top and Bottom are plateaus in the units of the Y axis.

**<u>Homogenous Time Resolved Fluorescence (HTRF) probe displacement assay</u>**

Assays used a buffer composition of 50 mM Tris pH 7.5, 150 mM NaCl, 2 mM $MgCl_2$, 0.01% w/v Triton X100 and 1 mM DTT. In screening mode to identify eIF2B ISRIB competitors, the HTRF assay was set up with a top concentration of 1 mM, with 1:3 dilutions over 12 points. Concentrations of biotinylated eIF2B, streptavidin-Tb cryptate and ISRIB-Alexa647 probe remained fixed at 2, 1 and 15 nM respectively and the assays normalised to a final DMSO concentration of 1%. Assays were incubated for 60 minutes and measured as endpoints on a BMG Labtech PHERAstar FSX using a 337/620/665 nm optics module and calculating the HTRF ratio of acceptor fluorescence divided by donor fluorescence. For elucidation of compound mechanism of action, probe displacement was monitored in matrix experiments as a function of both Compound A enantiomers and eIF2(αP) concentrations. eIF2(αP) was tested at 3, 1.5, 0.75, 0.38, 0.19, 0.094, 0.047 and zero μM, whilst the compound was tested with a top concentration of 1 mM, with 1:3 dilutions over 12 points. Data was fitted using GraphPad Prism 8 to the Yonetani-Theorell equation,

$$Y = \frac{Ymax}{1+\frac{[L]f}{Kf}+\frac{[L]v}{Kv}+\frac{[L]f[L]v}{\alpha Kf Kv}} + C$$

Where Y is the measured HTRF signal, $Y_{max}$ is the maximal HTRF signal, $[L]_f$ and $[L]_v$ are the concentrations of the fixed and varied ligands respectively, $K_f$ and $K_v$ are the apparent dissociation constants for the fixed and varied ligands respectively, α is a measure of the cooperativity of binding between the two ligands and C is the background HTRF signal.

**<u>Fluorescence Intensity (FI) - Guanine Exchange Factor (GEF) activity assay</u>**

Assays used a buffer composition of 20 mM HEPES pH 7.5, 120 mM KCl, 5 mM $MgCl_2$, 0.1% w/v bovine serum albumin (BSA) and 1 mM TCEP. Decrease in FI signal was monitored over time as a measure of GDP release, a surrogate for GEF activity[15]. All compounds were tested at a fixed or top concentration of 1 μM for 1:2 dilutions over 16 points. Concentrations of eIF2B, eIF2 (pre-loaded with GDP conjugated to BODIPY), and eIF2(αP) remained fixed at 10, 25 and 15.63 nM respectively. All assays were normalised to a final DMSO concentration of 1% and measured on a BMG Labtech PHERAstar FSX using a "FI 485 520" optics module. Data was fitted using GraphPad Prism 9 to the one-phase decay equation to determine half-lives of the dissociation curve:

$Y=(Y_0 - Plateau)*exp(-K*X) + Plateau$

Where Y is the measured FI signal, $Y_0$ is the maximal FI signal, X is the time in minutes, K is the rate constant equal to the reciprocal of X. $Y_0$ and Plateau are the same units as Y.

### Cryo-EM Methods

#### eIF2B:(P)eIF2α-NTD:Compound A-(*S*) cryo-EM grid preparation

2.5 μl of eIF2B:(P)eIF2α-NTD:Compound A-(*S*) complex purified by SEC was applied to a Quantifoil 1.2/1.3 Cu 300 mesh grid with 2 nm continuous carbon at 0.1 mg/ml final concentration. A Vitrobot Mark IV was used to vitrify the sample with blotting for 3s before plunging into liquid ethane.

Data was collected using a Krios G2 at the Cambridge Pharmaceutical Cryo-EM Consortium equipped with a Falcon IVi detector. 22,698 movies were collected at a nominal magnification of 120kx, pixel size 0.65 Å/pixel. A total dose of 60 $e^-/Å^2$ was applied over an exposure time of 4.25 s. A defocus range of -2.4 to -0.8 in 0.2 increments was used.

#### eIF2B:(P)eIF2α-NTD (apo) cryo-EM grid preparation

2.5 μl of eIF2B:(P)eIF2α-NTD mixed at a 1:4 ratio and incubated for 1 hour prior to grid freezing was applied to a glow-discharged Quantifoil 1.2/1.3 Cu 200 mesh grid. A Vitrobot Mark IV was used to vitrify the sample with blotting for 3 s before plunging into liquid ethane.

Data was collected using a Krios G4 equipped with a Falcon IVi detector. 6,728 moves were collected at a nominal magnification of 120kx, pixel size 0.64 Å/pixel. A total dose of 50 $e^-/Å^2$ was applied over an exposure time of 4.13 s. A defocus range of -2.4 to -0.8 in 0.2 increments was used.

#### Cryo-EM data processing

Images had been saved in EER format and upon import into cryoSPARC[29] were fractionated into 40 frames without upsampling. Patch motion correction and patch CTF correction were performed in cryoSPARC. Particles were initially picked using the blob picker, and 2D classes of the eIF2B complexes were then used to repick with template-based picking. Initial 2D classification of particles from the eIF2B:(P)eIF2α-NTD:Compound A-(*S*) dataset revealed multiple views of the eIF2B:(P)eIF2α-NTD complex but also of two contaminants the GroEL chaperone and a ribosome. The classes of these particles were removed and only those of eIF2B:(P)eIF2α-NTD taken forward into ab-initio reconstruction with just 1 class (291,184 particles). Homogeneous refinement followed this with defocus and CTF refinement performed on-the-fly. This resulted in an initial reconstruction at 2.67 Å overall resolution. 3D variability analysis in cryoSPARC[30] with 3 modes enabled separation of a subset of particles showing increased (P)eIF2α-NTD density quality, likely because of higher occupancy. Homogeneous refinement on this particle subset resulted in a reconstruction at 2.78 Å overall resolution.

Particles from the eIF2B:(P)eIF2α-NTD (apo) dataset again were initially picked using the blob picker and 2D classes showing secondary structure details were used for template-based picking. Ab-initio reconstruction with 1 class (120,653 particles) followed by homogeneous refinement with defocus and CTF refinement performed on-the-fly resulted in a reconstruction at 2.81 Å overall resolution. 3D variability analysis was also performed on the particles from this final reconstruction to compare the dynamics of the complex with and without Compound A-(*S*) present.

The .mrc map was converted to structure factors (.mtz) using CCPEM MRC to MTZ then viewed in WinCoot 0.8.9.1[31].

The SMILES string of the Compound A-(*S*) enantiomer was used to generate ligand dictionary (.cif) and coordinate (.pdb) files with grade from Global Phasing Ltd[32].

The PDB model of eIF2B:eIF2(αP) trimer 6K72 was used for model building into the eIF2B:(P)eIF2α:Compound A-(*S*) map. Model building was performed manually in Coot[31] using real-space refinement then the model was refined in Phenix[33].

Images were generated in UCSF Chimera[34] and Pymol[35].

<u>3D variability analysis on cryo-EM datasets</u>

Due to the differences in sample preparation methods used to obtain eIF2B:(P)eIF2α-NTD:Compound-A-(*S*) and eIF2B:(P)eIF2α-NTD (apo) datasets, 3D variability analysis was performed on both with a mask around just the eIF2B decamer.

### Cell culturing

U-2 OS cells (ATCC HTB-96) were cultured in growth media (DMEM (Gibco 41966-029) with 10% FBS (Gibco 011-90015M) and 1% P/S (Gibco 15140-122) in T175 flasks (Thermo Scientific 178885) until >70% confluency at 37 °C and 5% CO2. U-2 OS cells were used in the experiments described in Figure 4.

Adherent Chinese Hamster Ovary (CHO-K1) cells (ATCC CCL-61) were cultured in Ham's F12 nutrient mixture (Sigma), supplemented with 10% (v/v) serum (FetalClone-2, HyClone), 2 mM L-glutamine (Sigma), and 1% penicillin/streptomycin (Sigma). HEK293T cells (ATCC CRL-3216) were grown in DMEM media (Sigma) with the same supplements.

HeLa cervical epithelial cells (female) were cultured in DMEM high glucose medium (Sigma, D6546) supplemented with 10% Fetal Calf serum (FetalClone II, Thermo), 2 mM L- glutamine (G7513, Sigma Aldrich), 1x Penicillin/Streptomycin (P0781, Sigma), 1x non-essential amino acid solution (M7145, Sigma), and 55 μM β-mercaptothanol (Sigma, M3148). All cell types were incubated at 37 °C in a humidified environment containing 5% CO2. CHO cells were used in the experiments described in Figs. 4, 5 and 6 and Extended Data Figs. 4 and 5. As specified, cells were exposed to Compound B, Compound A-*(S)* or thapsigargin (Tg) for varying times and concentrations. These compounds were serially diluted in DMSO as necessary. Control cells were treated with DMSO alone.

### ATF4-luciferase assay

HEK293 cells were modified to express a modified version of a ATF4-luciferase reporter from Sidrauski et al. (2013)[12] containing a de-stabilised version of nanoluciferase (Promega) in place of the Firefly luciferase. Single cell clones were produced and the final clone chosen based upon optimal assay performance. Prior to assay, cells were grown to 70% confluency in selection media comprising DMEM (Sigma #D1145), FBS (Gibco 10% final), Glutamax (Thermofisher 1X final) and 0.5 μg/ml puromycin (Gibco). Cells were removed from culture flasks using TrypLE (Thermofisher) and diluted to 65000 cells/mL in medium. Tunicamycin was added to 1 μM final concentration to elicit the ISR. The falcon tube containing the cell suspension was then sealed and inverted 10 times to allow even distribution of tunicamycin. Cells were dispensed in a 40 μl volume into 384-well plates (Greiner 781080) using a Multidrop Combi (Thermofisher), pulse spun in a centrifuge at 100 x r.c.f. and then placed into an incubator. At 6 hours following plate dispense, media was removed from the plates using a gentle spin protocol on the bluewasher plate washer (Bluecatbio) and immediately replaced with 20 μl of a lysis solution comprising 1 part PBS + 1 part Nanoluc lysis buffer and the Nanoluc substrate at 1:100 dilution. Plates were pulse spun at 100 x r.c.f. and then incubated for 15 minutes at room temperature. Plates were then read for luminescence on the Envision plate reader (Perkin Elmer) with luminescence filter 700 (212) and luminescence mirror 404 and a 1 second measurement time.

**Stress Granule Formation Assay**

The U-2 OS cells were lifted using accutase (Gibco A11105-01) for 10 minutes at 37 °C and 5% CO2, spun down in growth media at 2000 RPM for 2 minutes, re-suspended and seeded at a density of 5,000 cells/well on Greiner 96-well microplates (655946) in 100ul growth media. The cells were cultured for 48 hours at 37 °C and 5% CO2. The U-2 OS cells should have flattened in morphology after 48 hours to allow for a large cytoplasmic surface area for stress granule formation and imaging.

At the time of treatment, Tg (Cayman Chemical Company 10522) and Compound A were prepared at 2x the concentration required in growth media and added at 50ul each (or DMSO vehicle control) for a total of 100 μl/well for 60 minutes at 37 °C and 5% $CO_2$. Cells were then fixed with 4% formaldehyde (Thermo Scientific 28906, diluted in DPBS (Gibco 14190-094)) for 15 minutes at RT, and free aldehydes quenched with 50 mM Ammonium Chloride (Sigma 09718-250G, dissolved in DPBS).

The cells were permeabilised with 0.1% Triton X-100 solution (Sigma 102378814, diluted in DPBS) for 10 minutes at RT and blocked with 2% BSA (Sigma A9576, diluted in DPBS) for 60 minutes at RT. The primary antibody solution (1% BSA in DPBS with 1:200 G3BP1 antibody (mouse, BD Biosciences 611127)) was applied for 60 minutes at RT or overnight at 4 °C prior to thrice DPBS washes and secondary antibody solution (1% BSA in DPBS with 1:1,000 AlexaFluor-647 goat anti-mouse (Thermo Scientific A21235) and 1:10,000 Hoechst (Thermo Scientific H3570)) addition for 60 minutes at RT in the dark. After thrice DPBS wash, the plate was sealed with a silver foil plate seal and imaged on the Molecular Devices ImageXpress at 60X magnification.

The images (cell boundaries, cell nuclei and cytoplasm, and stress granules) were segmented and quantified on the Molecular Devices MetaXpress software, exported to Excel, and the percentage of stress granule-positive cells calculated and analysed on Excel and GraphPad Prism 9.0.

**CHOP::GFP and Luciferase reporter assay**

Engineering of CHO-C30 containing the CHOP::GFP and CHO-S21 cells containing dual reporter, CHOP::GFP and XBP1::Turquoise, was described previously[36, 37].

CHO-C30 CHOP::GFP cells containing either endogenous eIF2B$\delta^{L180F}$ (clone S7) or eIF2B$\beta^{H188K}$ (clone X) was reported in the past [13, 14].

CHO-S21 CHOP::GFP and Xbp1::Turquoise dual reporter cells bearing non-phosphorylatable eIF2$\alpha^{S51A}$ was documented prior[37].

Treatment of the reporter cells with a chemical stressor L-histidinol (ACROS Organics) increases the amount of uncharged histidyl-tRNAs that induces one of the eIF2α kinases, GCN2. eIF2(αP) inhibits the GEF eIF2B, initiating the ISR and activating CHOP::GFP, which is detected by fluorescent activated cell sorting (FACS).

CHO CHOP::GFP reporter cells were plated onto 12-well plate at the density of 30 x $10^3$ cells/well 48 hours prior the treatment. Treatment with chemical compounds was performed in fresh media between 20-24 hours or as indicated. Concentrations of the drugs used for each experiment are indicated in figures or figure legends. Cells were then washed twice with ice-cold PBS and collected in ice-cold PBS supplemented with 4 mM EDTA. Single cell fluorescent signals (10,000/sample) was measured by FACS Calibur (Beckton Dickinson). Data analysed with FlowJo software (BD Life Sciences).

The CHOP:: Luciferase assay was performed by following the previously reported methods (Harding et al., 2005). CHO-KI cells stably expressing CHOP::luciferase reporter gene were plated at a density of 2 x $10^4$ cells per well in 96-well plates 24 hr before adding fresh media containing 10 μM of compounds in DMSO. Sixteen hours later, cells were washed twice and lysed in 25 μl lysis buffer (25 mM glygly, 15 mM $MgSO_4$, 4 mM EGTA, 1% Triton X-100, and 1 mM DTT [added fresh]), and luciferase activity was measured following the addition of 25 μl assay buffer (25 mM glygly, 15 mM $MgSO_4$, 4 mM EGTA, 1% Triton X-100, 11.7 mM potassium phosphate, 1.6 mM ATP, 0.2 mg/ml coenzyme A, 500 μM luciferin, and 2 mM DTT). The Luminescence were detected by using a CLARIOstar plate reader.

**Polysomes profiling**

Polysome analyses were carried out as reported previously[4]. HEK cells over 90% confluence were treated for 1 hr with Compound B (30 μM) or Tg (1 μM). Cycloheximide (0.1 mg/ml) was added during the last 5 min of treatment. The plates were then washed twice with 5 ml ice-cold PBS containing 0.1 mg/ml cycloheximide, and cells were collected by scraping into 1 ml of PBS. The cells were lysed in 350 ⌠l polysome extraction buffer (15 mM Tris-Cl [pH 7.4], 15 mM $MgCl_2$, 0.3 M NaCl, 1% Triton X-

100, 0.1 mg/ml cycloheximide, 1 mg/ml heparin, 1 mM phenylmethysulfonyl fluoride, 4 μg/ml aprotonin, and 2 μg/ml pepstatin A) and incubated on ice for 10 min. The lysates were cleared by centrifugation at 15,000 rpm and layered onto a 5mL 10%–60% sucrose gradient prepared in the extraction buffer, and, after a 1 hr spin at 45000 rpm in an SW55 Ti rotor, the absorbance across the gradient at 260 nm analysed by using a Biocomp Piston Gradient Fractionator™ with the Triax™ FC-2-UV flow cell detector.

**Flow Cytometry Analysis**

To assess ISR induction, cells were cultured in 12-well plates for 40-48 hours before being treated with the specified compounds for the indicated period. For flow cytometry analysis, the cells were washed twice with PBS and then collected in PBS containing 4 mM EDTA. A total of 20,000 cells per sample were analysed using a multi-channel flow cytometer, the LSR Fortessa cell analyser (BD Biosciences). Live cells were gated based on FSC-A/SSC-A parameters, and singlets were gated based on FSC-W/SSC-A parameters. CHOP: GFP fluorescence signals were detected using a 488 nm excitation laser and a 530/30 nm emission filter. FlowJo V10 was used for data analysis, and median reporter analysis was performed using Prism 10 (GraphPad).

**RNA isolation and reverse transcription**

To isolate total RNA, cells were treated with TRIzol™ Reagent (Invitrogen) for 10 minutes and subsequently transferred to fresh tubes. After addition of 200 μl of chloroform, the samples were vortexed for 1 minute and then centrifuged at 13,500 x g for 15 minutes at 4°C. The upper aqueous layer was collected and mixed with an equal volume of 70% ethanol. The samples were applied to PureLink™ RNA Mini Kit columns and purified and eluted as per the manufacturer's instructions (Invitrogen). RNA concentration was determined using a NanoDrop, and 2 μg of RNA from each sample was assessed for integrity by electrophoresis on a 1.2% agarose gel.

For reverse transcription (RT)-PCR, 2 μg of RNA was first heated at 70 °C for 10 minutes in RT buffer (Thermo Scientific; Cat #EP0441) containing 0.5 mM dNTP and 0.05 mM Oligo dT18. The reaction was then supplemented with 0.5 μl of RevertAid Reverse Transcriptase (Thermo Scientific) and 100 mM DTT, followed by incubation at 42 °C for 90 minutes. The resulting cDNA was diluted up to 1:4.

**PCR analysis of XBP1 mRNA splicing**

XBP1s and XBP1u fragments were amplified from cDNA via PCR using primers (hamXBP1.19S: GGCCTTGTAATTGAGAACCAGGAG, mXBP1.14AS: GAATGCCCAAAAGGATATCAGACTC) flanking the IRE1-identified splicing site, with the NEB Q5® High-Fidelity 2X Master Mix, by following the method reported by Tung et al. (2024). Briefly, XBP1u (255 bp) and XBP1s (229 bp) were resolved on a 3% agarose gel through electrophoresis and stained with SYBR Safe nucleic acid gel stain (Invitrogen). A hybrid XBP1 band (280 bp) was also observed. The XBP1s percentage was determined by analysing band intensity with Fiji, v1.53c, assuming that 50% of the XBP1 hybrid signal represented XBP1s and the other 50% represented XBP1u.

**[35S] metabolic labelling**

The [35S] metabolic labelling was conducted as previously reported procedure[38]. CHO-K1 cells were treated with Compound B or Tg at specified concentrations for 30 minutes, followed by a 20 minute starvation period in methionine/cysteine-free DMEM (21013024, GIBCO) in the presence of Compound B or Tg. Subsequently, the cells were pulsed for 10 minutes with 5.5 µCi/well [35S] methionine/cysteine (Expre35S Protein Labeling Mix) before being harvested. After the pulse, the culture media were collected, and the cells were harvested on ice in PBS supplemented with cycloheximide (0.1 mg/ml). Cell pellets were collected by centrifugation at 3000 rpm for 5 minutes at 4 °C. The cells were then lysed using Nonidet lysis buffer (150 mM NaCl, 50 mM Tris-HCl pH 7.5, 1% (v/v) NP-40, 0.05 mM TCEP) supplemented with cycloheximide (0.1 mg/ml), 1 mM phenylmethylsulfonyl fluoride, 4 µg/ml aprotinin, and 2 µg/ml pepstatin A. A post-nuclear supernatant was prepared by centrifugation at 20,000 × g at 4°C for 15 minutes. Radiolabeled cell lysates were mixed with 4 × SDS-PAGE loading buffer, incubated at 55 °C for 10 minutes, separated on 12.5% SDS-PAGE gels, and detected by autoradiography using a Typhoon biomolecular imager (GE Healthcare). Quantification was performed using Fiji (ImageJ).

| | #1 eIF2B + (P)eIF2α-NTD + Compound A-*(S)* (EMDB-xxxx) (PDB xxxx) |
|---|---|
| **Data collection and processing** | |
| Magnification | |
| Voltage (kV) | 300 |
| Electron exposure (e–/Å$^2$) | 40 |
| Defocus range (μm) | 0.8-2.4 |
| Pixel size (Å) | 0.65 |
| Symmetry imposed | C1 |
| Initial particle images (no.) | 2,096,572 |
| Final particle images (no.) | 125,874 |
| Map resolution (Å) | 2.78 |
| FSC threshold=0.143 | |
| **Refinement** | |
| Initial model used (PDB code) | 6K72 |
| Model resolution (Å) | 3.1 |
| FSC threshold=0.5 | |
| Map sharpening *B* factor (Å$^2$) | -53.7 |
| Model composition | |
| Non-hydrogen atoms | 24,436 |
| Protein residues | 3,404 |
| Ligands | 77 |
| *B* factors (Å$^2$) | |
| Protein | 124.9 |
| Ligand | 92.5 |
| R.m.s. deviations | |
| Bond lengths (Å) | 0.003 |
| Bond angles (°) | 0.517 |
| Validation | |
| MolProbity score | 1.92 |
| Clashscore | 4.26 |
| Poor rotamers (%) | 4.2 |
| Ramachandran plot | |
| Favored (%) | 96.26 |
| Allowed (%) | 3.74 |
| Disallowed (%) | 0.00 |

**Supplementary Table 1: Cryo-EM data collection, refinement and validation statistics**

### Chemistry Experimental

#### Abbreviations

Ac, acetyl; ACN, acetonitrile; DIPEA, *N,N*-diisopropylethylamine; DMF, *N,N*-dimethylformamide; DCM, dichloromethane; DMSO, *N,N*-dimethylsulfoxide; EA, ethyl acetate; Et, ethyl; FA, formic acid; Me, methyl; PE, petroleum ether; rt, room temperature; TFA, trifluoroacetic acid; TEA, triethylamine.

#### Chemistry General Experimental

Reagents and solvents were obtained from commercial suppliers and used without any further purification unless otherwise stated. All reagents were weighed and handled in air unless otherwise stated. Purity and characterization of compounds were established by a combination of liquid chromatography-mass spectroscopy (LC-MS) and NMR analytical techniques and was >95% for all test compounds.

#### NMR Spectroscopy

$^{1}$H and $^{13}$C NMR spectra were recorded using Bruker Avance III, Avance III HD or Avance III NEO spectrometers at a proton frequency of 300 MHz, or a Bruker Avance Neo spectrometer at a proton frequency of 500 MHz and were determined in $CDCl_3$, DMSO-$d_6$ or MeOH-$d_4$. $^{19}$F NMR spectra are reported in ppm from $CFCl_3$ and are uncorrected. $^{1}$H NMR chemical shifts ($\delta$) are reported in ppm ± 0.01 relative to TMS (0.00 ppm) or solvent peaks as the internal reference. Splitting patterns are indicated as follows: s, singlet; d, doublet; t, triplet; q, quartet; m, multiplet; br, broad peak. Coupling constants ($J$) are given in Hertz (Hz) ± 0.1 Hz. $^{13}$C NMR and $^{19}$F chemical shifts (δ) are given in ppm ±0.1 except in those cases whereby rounding to ±0.1 would result in the signals having the same absolute value.

#### Mass Spectrometry

LC/MS analyses were performed using a Shimadzu LCMS-2020 with electrospray ionization in positive ion detection mode with 20ADXR pump, SIL-20ACXR autosampler, CTO-20AC column oven, M20A PDA Detector and LCMS 2020 MS detector. LC was run in two set ups: 1) Halo C18 column (2.0 µm 3.0x30 mm) in combination with a gradient (5-100% B in 1.2 minutes) of water and FA (0.1%) (A) and $CH_3CN$ and FA (0.1%) (B) at a flow rate of 1.5 mL/min; 2) Poroshell HPH C18 column (2.7 µm 3.0x50 mm) in combination with a gradient (5-95% B in 2 minutes) of aqueous 46 mM ammonium carbonate/ammonia buffer at pH 10 (A) and MeCN (B) at a flow rate of 1.2 mL/min ; 3) Halo C18 column (2.0 µm 3.0x30 mm) in combination with a gradient (5-95% B in 2 minutes) of water and TFA (0.05%) (A) and $CH_3CN$ and TFA (0.05%) at a flow rate of 1.5 mL/min (B). The Column Oven (CTO-20AC) temperature was 40.0°C. The injection volume was 1 µl. PDA (SPD-M20A) detection was in the range 190–400 nm. The MS detector, which was configured with electrospray ionization as ionizable source; Acquisition mode: Scan; Nebulizing Gas Flow:1.5 L/min; Drying Gas Flow:15 L/min; Detector Voltage: Tuning Voltage ± 0.2 kv; DL Temperature: 250 ºC; Heat Block Temperature: 250 ºC; Scan Range: 90.00 - 900.00 m/z.

#### High-Resolution Mass Spectrometry

Accurate mass data of samples were obtained using HRMS system Thermo Orbitrap IQ-X. The samples were separated on reversed phase Acquity UPLC CSH C18, 2.1mm × 50mm, 1.7 $\mu$m particles using gradient elution with 5% $CH_3CN$ to 95% $CH_3CN$ in 3.5 minutes.; 0.1% Formic acid, pH2/5% $CH_3CN$ to 95% $CH_3CN$ in 4 minutes or 0.1% Ammonium hydroxide, pH 10. Acquity PDA Detector was in the range 220–400 nm. The injection volume 1.0 $\mu$L. Inlet system: Thermo Vanquish autosampler, PDA detector and binary pump. The column temperature was set at 45 °C and the flow rate at 0.75 mL/min. Instrument control and accurate mass data were processed using Xcalibur software. The MS data were acquired in positive and negative ionization mode with the following conditions: MS equipped with HESI with capillary voltage at 3.0 kV (+) and 2.5kV (-), ion transfer temp at 350 °C, source temperature at 350 °C, Aux gas flow = 15, sheath gas flow = 60 and sweep gas = 2 (all arbitrary units), acquisition mass range of 100 – 1000 Da.

**Flash Column Chromatography**
Chromatographic purification of products was accomplished with a CHEETAH® MP200 system with integrated UV detection, using column chromatography on (A) silica gels (40-60 µm), eluted with petroleum ether and ethyl acetate or dichloromethane and methanol, (B) C18 spherical (20-35 µm), eluted with water and acetonitrile or MeOH with formic acid (0.1%) or NH4HCO3 (10 mmol) modifier.

**Preparative HPLC**
Preparative HPLC was performed with a Waters MassLynx system with integrated MS detection and equipped with Prep C18 OBD 5µm 30 x 150 mm columns from XBridge or Xselect CSH. Alternatively, Gilson GX-281 with integrated UV detection was used, equipped with either XBridge or Sunfire C18 10µm, 19x150 ID or 19x250mm. As eluent (acidic) gradients of water/MeCN/FA acid (95/5/0.1) or water/0.05% TFA (A) and MeCN/ MeOH (B) or 0.05% ammonia in water /10 mmol NH4HCO3 (A) and MeCN/ MeOH (B) were applied.

**Preparative Chiral-HPLC**
Preparative Chiral HPLC was performed with a Gilson GX-281 system with integrated UV detection and equipped with one of Chiralpak AS, AD, Chiralcel OD,OJ Chiralpak IA,IB,IC,ID,IE,IF,IG,IH columns (Daicel Chemical Industries, Ltd.) (R,R)-Whelk-O1, (S,S)-Whelk-O1 columns (Regis technologies, Inc. ) CHIRAL Cellulose-SB, SC, SA columns (YMC Co., Ltd.) at different column size (250x20mm, 250x30mm) with noted percentage of either ethanol in hexane (%Et/Hex) or isopropanol in hexane (%IPA/Hex) as isocratic solvent systems.

**Synthesis of Compound A**

**Scheme 1**

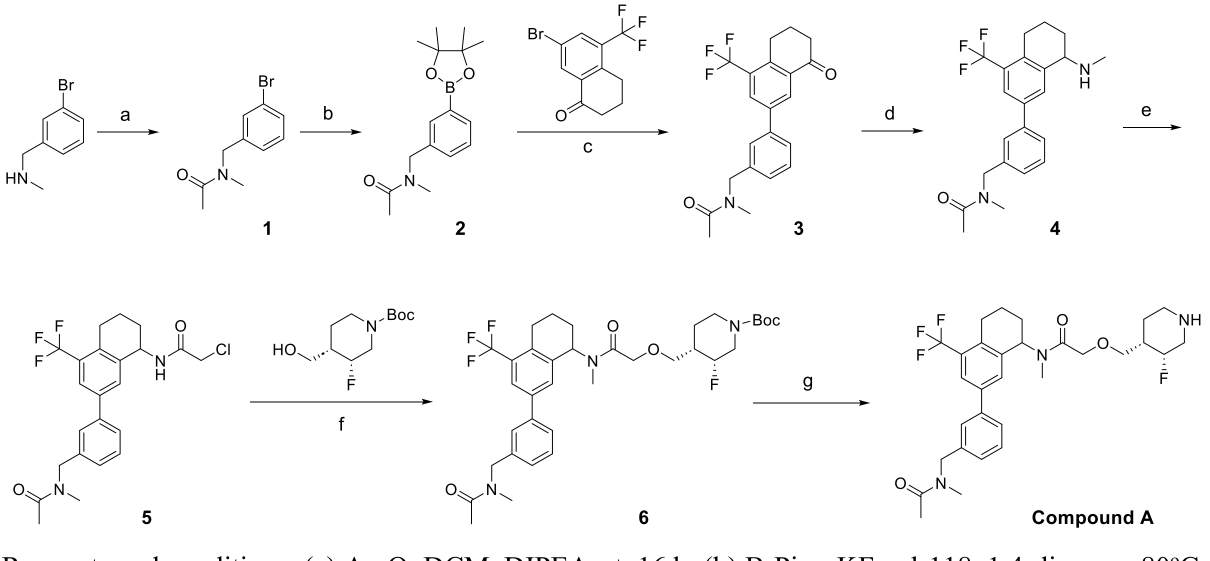

Reagents and conditions: (a) $Ac_2O$, DCM, DIPEA, rt, 16 h; (b) $B_2Pin_2$, KF, pd-118, 1,4-dioxane, 80ºC, 16 h; (c) XPhos Pd G3, $Cs_2CO_3$, 1,4-dioxane, $H_2O$, 100ºC, 2 h; (d) $MeNH_2HCl$, $NaBH_3CN$, DIPEA, EtOH, rt, 2 h; (e) 2-chloroacetyl chloride, DCM, DIPEA, rt, 2 h; (f) NaH, THF, rt, 16 h; (g) 4M HCl in 1,4-dioxane, DCM, rt, 1 h

**N-(3-bromobenzyl)-N-methylacetamide (1)**

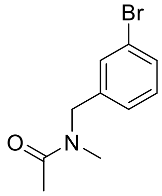

To a stirred solution of [(3-bromophenyl)methyl](methyl)amine (10 g, 49.98 mmol, 1 equiv) and $Ac_2O$ (10.20 g, 99.96 mmol, 2 equiv) in DCM (100 mL) was added DIPEA (19.38 g, 149.94 mmol, 3 equiv) at room temperature. The resulting mixture was stirred for 16 h at room temperature.

The reaction was quenched by the addition of water (200 mL). The resulting mixture was extracted with DCM (3 x 100 mL). The combined organic layers were washed with brine (1 x 100 mL), dried over anhydrous $Na_2SO_4$. After filtration, the filtrate was concentrated under reduced pressure.

The residue was purified by silica gel column chromatography, elution gradient 0 to 100% EA in PE. Pure fractions were evaporated to afford N-[(3-bromophenyl)methyl]-N-methylacetamide (**1**) (11 g, 91%) as a white solid.

MS (ESI) *m/z* calcd for $C_{10}H_{12}BrNO$ $[M+H]^+$: 242; found: 242.

**N-methyl-N-(3-(4,4,5,5-tetramethyl-1,3,2-dioxaborolan-2-yl)benzyl)acetamide (2)**

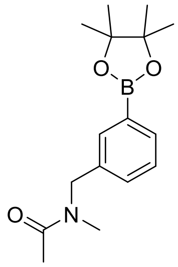

To a stirred solution of N-[(3-bromophenyl)methyl]-N-methylacetamide (5 g, 20.65 mmol, 1 equiv) and KF (3.60 g, 61.95 mmol, 3 equiv) in dioxane (100 mL) were added $PdCl_2$(dtbpf) (1.35 g, 2.06 mmol, 0.1 equiv) and 4,4,5,5-tetramethyl-2-(tetramethyl-1,3,2-dioxaborolan-2-yl)-1,3,2-dioxaborolane (15.73 g, 61.95 mmol, 3 equiv). The resulting mixture was stirred for 16 h at 80°C under nitrogen atmosphere.

The resulting mixture was concentrated under reduced pressure. The residue was dissolved in water (200 mL) and extracted with EtOAc (3 x 100 mL). The combined organic layers were washed with brine (1 x 100 mL), dried over anhydrous $Na_2SO_4$. After filtration, the filtrate was concentrated under reduced pressure.

The residue was purified by silica gel column chromatography, elution gradient 0 to 100% EA in PE. Pure fractions were evaporated to afford 3-[(N-methylacetamido)methyl]phenylboronic acid (**2**) (3.2 g, 75%) as a white solid.

MS (ESI) *m/z* calcd for $C_{16}H_{24}BNO_3$ $[M+H]^+$: 290; found: 290.

**N-methyl-N-(3-(8-oxo-4-(trifluoromethyl)-5,6,7,8-tetrahydronaphthalen-2-yl)benzyl)acetamide (3)**

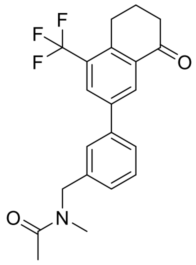

To a stirred solution of N-methyl-N-{[3-(4,4,5,5-tetramethyl-1,3,2-dioxaborolan-2-yl)phenyl]methyl}acetamide (1 g, 3.46 mmol, 1 equiv) and 7-bromo-5-(trifluoromethyl)-3,4-dihydro-2H-naphthalen-1-one (1.11 g, 3.80 mmol, 1.1 equiv) in dioxane (16 mL) and $H_2O$ (4 mL) were added XPhos Pd G3 (0.29 g, 0.35 mmol, 0.1 equiv) and $Cs_2CO_3$ (2.82 g, 8.65 mmol, 2.5 equiv). The resulting mixture was stirred for 2 h at 100°C under nitrogen atmosphere.

The resulting mixture was concentrated under reduced pressure. The residue was dissolved in water (200 mL) and extracted with EtOAc (3 x 100 mL). The combined organic layers were washed with brine (1 x 100 mL), dried over anhydrous $Na_2SO_4$. After filtration, the filtrate was concentrated under reduced pressure.

The residue was purified by silica gel column chromatography, elution gradient 0 to 100% EA in PE. Pure fractions were evaporated to afford N-methyl-N-({3-[8-oxo-4-(trifluoromethyl)-6,7-dihydro-5H-naphthalen-2-yl]phenyl}methyl)acetamide (**3**) (350 mg, 27%) as a white solid.

MS (ESI) *m/z* calcd for $C_{21}H_{20}F_3NO_2$ $[M+H]^+$: 376; found: 376; $^1H$ NMR (300 MHz, DMSO-*d6*) δ 2.08 (s, 3H), 2.08-2.16 (m, 2H), 2.71-2.75 (m, 2H), 2.89 (s, 3H), 3.11-3.14 (m, 2H), 4.59 (s, 1H), 4.67 (s, 1H), 7.25-7.30 (m, 1H), 7.45–7.72 (m, 3H), 8.13–8.2 (m, 1H), 8.37-8.39 (m, 1H).

**_Rac_-N-methyl-N-(3-(8-(methylamino)-4-(trifluoromethyl)-5,6,7,8-tetrahydronaphthalen-2-yl)benzyl)acetamide (4)**

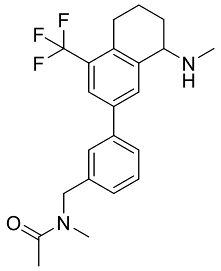

To a stirred solution of N-methyl-N-({3-[8-oxo-4-(trifluoromethyl)-6,7-dihydro-5H-naphthalen-2-yl]phenyl}methyl)acetamide (340 mg, 0.91 mmol, 1 equiv) and $CH_3NH_2HCl$ (306 mg, 4.53 mmol, 5 equiv) in EtOH (10 mL) were added NaBH3CN (285 mg, 4.53 mmol, 5 equiv) and DIPEA (585 mg, 4.53 mmol, 5 equiv). The resulting mixture was stirred for 2h at room temperature.

The resulting mixture was concentrated under reduced pressure. The residue was dissolved in water (50 mL) and extracted with EtOAc (3 x 50 mL). The combined organic layers were washed with brine (1 x 50 mL), dried over anhydrous $Na_2SO_4$. After filtration, the filtrate was concentrated under reduced pressure.

The residue was purified by reversed-phase flash chromatography with the following conditions: column, C18 silica gel; mobile phase, MeCN in Water (0.1% FA), 0% to 100% gradient in 40 min; detector, UV 220 nm. This resulted in N-methyl-N-({3-[8-(methylamino)-4-(trifluoromethyl)-5,6,7,8-tetrahydronaphthalen-2-yl]phenyl}methyl)acetamide (**4**) (300 mg, 85%) as a white solid.

MS (ESI) *m/z* calcd for $C_{22}H_{25}F_3N_2O$ $[M+H]^+$: 391; found: 391.

***Rac*-2-chloro-N-(7-(3-((N-methylacetamido)methyl)phenyl)-5-(trifluoromethyl)-1,2,3,4-tetrahydronaphthalen-1-yl)acetamide (5)**

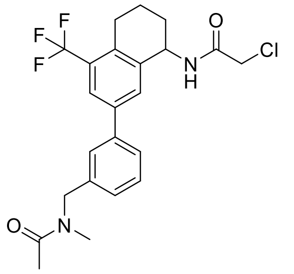

To a stirred solution of N-methyl-N-({3-[8-(methylamino)-4-(trifluoromethyl)-5,6,7,8-tetrahydronaphthalen-2-yl]phenyl}methyl)acetamide (290 mg, 0.74 mmol, 1 equiv) and DIPEA (0.26 mL, 1.49 mmol, 2 equiv) in DCM (10 mL) was added chloroacetyl chloride (0.08 mL, 0.67 mmol, 1 equiv) dropwise. The resulting mixture was stirred for 2 h at room temperature.

The reaction was quenched by the addition of water (50 mL) at room temperature. The resulting mixture was extracted with DCM (3 x 50 mL). The combined organic layers were washed with brine (1 x 50 mL), dried over anhydrous $Na_2SO_4$. After filtration, the filtrate was concentrated under reduced pressure. This resulted in 2-chloro-N-methyl-N-(7-{3-[(N-methylacetamido)methyl]phenyl}-5-(trifluoromethyl)-1,2,3,4-tetrahydronaphthalen-1-yl)acetamide (**5**) (320 mg, 92.27%) as a light yellow solid used in the next step directly without further purification.

**tert-butyl (3R,4S)-3-fluoro-4-((2-(methyl(7-(3-((N-methylacetamido)methyl)phenyl)-5-(trifluoromethyl)-1,2,3,4-tetrahydronaphthalen-1-yl)amino)-2-oxoethoxy)methyl)piperidine-1-carboxylate (6)**

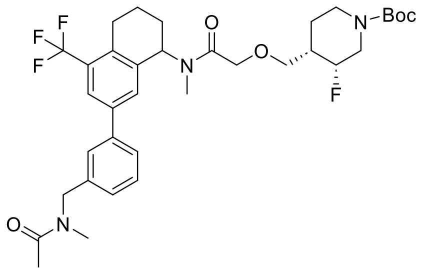

To a stirred solution of tert-butyl (3R,4S)-3-fluoro-4-(hydroxymethyl)piperidine-1-carboxylate (90 mg, 0.39 mmol, 1.2 equiv) in DMF (5 mL) was added NaH (15 mg, 0.39 mmol, 1.2 equiv, 60%). The resulting mixture was stirred for 0.5 h at room temperature under nitrogen atmosphere. Then 2-chloro-N-methyl-N-(7-{3-[(N-methylacetamido)methyl]phenyl}-5-(trifluoromethyl)-1,2,3,4-tetrahydronaphthalen-1-yl)acetamide (150 mg, 0.32 mmol, 1 equiv) was added into the above solution. The final reaction mixture was stirred for 16 h at room temperature.

The reaction was quenched by the addition of Water/Ice (20 mL). The resulting mixture was extracted with EtOAc (3 x 20 mL). The combined organic layers were washed with brine (1 x 20 mL), dried over anhydrous $Na_2SO_4$. After filtration, the filtrate was concentrated under reduced pressure.

The residue was purified by reverse flash chromatography with the following conditions: column, C18 silica gel; mobile phase, MeCN in water(0.1% TFA), 0% to 100% gradient in 30 min; detector, UV 254 nm. This resulted in tert-butyl (3R,4S)-3-fluoro-4-({[methyl(7-{3-[(N-methylacetamido)methyl]phenyl}-5-(trifluoromethyl)-1,2,3,4-tetrahydronaphthalen-1-yl)carbamoyl]methoxy}methyl)piperidine-1-carboxylate (**6**) (130 mg, 61%) as a white solid.

MS (ESI) *m/z* calcd for $C_{36}H_{45}F_4N_3O_5$ $[M+Na]^+$: 686; found: 686.

**2-(((3R,4S)-3-fluoropiperidin-4-yl)methoxy)-N-methyl-N-(7-(3-((N-methylacetamido)methyl)phenyl)-5-(trifluoromethyl)-1,2,3,4-tetrahydronaphthalen-1-yl)acetamide (Compound A)**

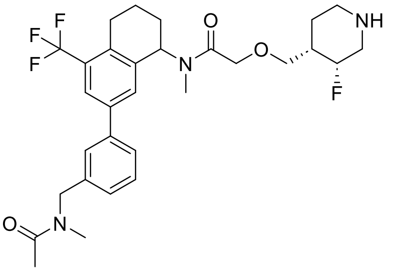

To a stirred solution of tert-butyl (3R,4S)-3-fluoro-4-({[methyl(7-{3-[(N-methylacetamido)methyl]phenyl}-5-(trifluoromethyl)-1,2,3,4-tetrahydronaphthalen-1-yl)carbamoyl]methoxy}methyl)piperidine-1-carboxylate (110 mg, 0.17 mmol, 1 equiv) in DCM (5 mL) was added dropwise 4M HCl in 1,4-dioxane (1 mL, 4 mmol/L, 24 equiv) at room temperature. The resulting mixture was stirred for 1 h at room temperature.

The resulting mixture was concentrated under reduced pressure, then diluted with DCM (20 mL), basified to pH 8 with saturated $NaHCO_3$ (aq.). The resulting mixture was extracted with DCM (3 x 20 mL). The combined organic layers were washed with brine (1 x 20 mL), dried over anhydrous $Na_2SO_4$. After filtration, the filtrate was concentrated under reduced pressure.

The residue was purified by reverse flash chromatography with the following conditions: column, C18 silica gel; mobile phase, MeCN in water(0.1% FA), 0% to 100% gradient in 20 min; detector, UV 220 nm. This resulted in 2-{[(3R,4S)-3-fluoropiperidin-4-yl]methoxy}-N-methyl-N-(7-{3-[(N-methylacetamido)methyl]phenyl}-5-(trifluoromethyl)-1,2,3,4-tetrahydronaphthalen-1-yl)acetamide (**Compound A**) (42 mg, 42%) as an off-white solid.

MS (ESI) *m/z* calcd for $C_{30}H_{37}F_4N_3O_3$ $[M+Na]^+$: 586; found: 586; HRMS (ESI) *m/z* calcd for $C_{30}H_{37}F_4N_3O_3$ $[M+H]^+$: 564.2849; found: 564.2857; $^1H$ NMR (300 MHz, MeOD) δ 1.60–2.37 (m, 7H), 2.18 (s, 3H), 2.62–2.81 (m, 3H), 2.95 (s, 2H), 3.04-3.29 (m, 4H), 3.34–3.77 (m, 4H), 4.22–4.6 (m, 2H), 4.6–4.78 (m, 2H), 4.90-5.31 (m, 2H), 5.89 (m, 1H), 7.27-7.29 (m, 1H), 7.36–7.61 (m, 4H), 7.73–7.84 (m, 1H). Assigned: 36, one hydrogen exchanged, $^{13}C$ NMR (125 MHz, $CDCl_3$) δ 21.5, 21.8, 24.5, 25.8, 26.3, 27.9, 29.2, 29.9, 33.8, 35.8, 39.8, 45.2, 49.7, 50.6, 52.9, 54.1, 56.8, 70.4, 71.5, 72.2, 85.8, 87.2, 122.9 - 130.0 (m), 136.5 - 140.8 (m), 169.6, 170.1, 170.8; $^{19}F$ NMR (471 MHz, $CDCl_3$) δ -61.14 (3F), -204.69 (1F).

**Synthesis of Compound A-(*S*) and A-(*R*)**

**Scheme 2**

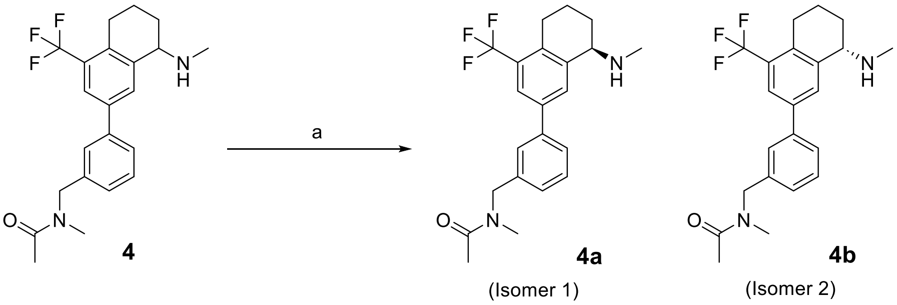

Reagents and conditions: (a) chiral separation

*rac*-N-methyl-N-(3-(8-(methylamino)-4-(trifluoromethyl)-5,6,7,8-tetrahydronaphthalen-2-yl)benzyl)acetamide (**4**) was purified by preparative chiral-HPLC on Column: CHIRALPAK ID-3, 4.6*50mm; 3um; Mobile Phase A: Hexane (0.1%DEA): EtOH=80: 20; Flow rate: 1 mL/min. The fractions containing the desired compound were evaporated to dryness to afford (*R*)-N-methyl-N-(3-(8-(methylamino)-4-(trifluoromethyl)-5,6,7,8-tetrahydronaphthalen-2-yl)benzyl)acetamide (270 mg, 31 %) as a colourless gum and (*S*)-N-methyl-N-(3-(8-(methylamino)-4-(trifluoromethyl)-5,6,7,8-tetrahydronaphthalen-2-yl)benzyl)acetamide (300 mg, 35 %) as a colourless gum. The stereochemistry was assigned using VCD.

**(*R*)-N-methyl-N-(3-(8-(methylamino)-4-(trifluoromethyl)-5,6,7,8-tetrahydronaphthalen-2-yl)benzyl)acetamide (4a)**

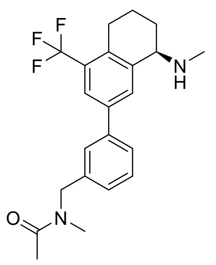

Chiral HPLC, Column: CHIRALPAK ID-3 4.6*50mm 3μm, Mobile phase: Hexane(0.1%DEA):EtOH=80:20, er = 98.6:1.4; MS (ESI) *m/z* calcd for $C_{22}H_{25}F_3N_2O$ $[M+H]^+$: 391; found: 391; $^1$H NMR (300 MHz, DMSO-*d6*) δ 1.59–1.75 (m, 1H), 1.75–1.86 (m, 2H), 1.87–2.01 (m, 1H), 2.06 (s, 3H), 2.11 (s, 1H), 2.33 (s, 3H), 2.71-2.90 (m, 2H), 2.99 (s, 3H), 3.68-3.75 (m, 1H), 4.59 (s, 1H), 4.66 (s, 1H), 7.18-7.24 (m, 1H), 7.39–7.65 (m, 3H), 7.7–7.96 (m, 2H); $^{19}$F NMR (300 MHz, DMSO-*d6*) δ -59.7.

**(S)-N-methyl-N-(3-(8-(methylamino)-4-(trifluoromethyl)-5,6,7,8-tetrahydronaphthalen-2-yl)benzyl)acetamide (4b)**

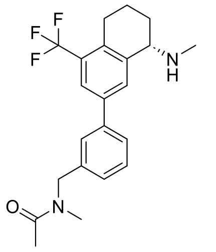

Chiral HPLC, Column: CHIRALPAK ID-3 4.6*50mm 3μm, Mobile phase: Hexane(0.1%DEA):EtOH=80:20, er = 99.8:0.2; MS (ESI) *m/z* calcd for $C_{22}H_{25}F_3N_2O$ $[M+H]^+$: 391; found: 391; $^1$H NMR (300 MHz, DMSO-*d6*) δ 1.62-1.75 (m, 1H), 1.75-1.86 (m, 2H), 1.87–2.01 (m, 1H), 2.06 (s, 3H), 2.06-2.21 (m, 1H), 2.33 (s, 3H), 2.70-2.92 (m, 2H), 2.93 (s, 3H), 3.70 (t, *J* = 5.1 Hz, 1H), 4.59 (s, 1H), 4.67 (s, 1H), 7.15-7.27 (m, 1H), 7.39–7.65 (m, 3H), 7.69–7.96 (m, 2H); $^{19}$F NMR (300 MHz, DMSO-*d6*) δ -59.7.

**2-(((3R,4S)-3-fluoropiperidin-4-yl)methoxy)-N-methyl-N-((R)-7-(3-((N-methylacetamido)methyl)phenyl)-5-(trifluoromethyl)-1,2,3,4-tetrahydronaphthalen-1-yl)acetamide (Compound A-(*R*))**

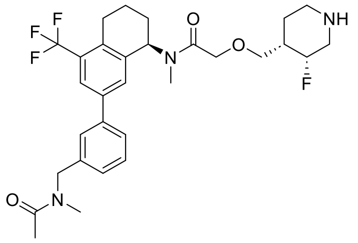

Prepared in an analogous method to **Compound A, Scheme 1,** where (*R*)-N-methyl-N-(3-(8-(methylamino)-4-(trifluoromethyl)-5,6,7,8-tetrahydronaphthalen-2-yl)benzyl)acetamide was used in place of *Rac*-N-methyl-N-(3-(8-(methylamino)-4-(trifluoromethyl)-5,6,7,8-tetrahydronaphthalen-2-yl)benzyl)acetamide. Resulting in the title compound (138 mg, 15%) as a white solid.

Chiral HPLC, Column: CHIRALCEL OD-3, Mobile Phase: Hexane(0.1%DEA):EtOH=85:15, dr = 99.7:0.3; MS (ESI) *m/z* calcd for $C_{30}H_{37}F_4N_3O_3$ $[M+H]^+$: 564.3; found: 564.4. HRMS (ESI) *m/z* calcd for $C_{30}H_{37}F_4N_3O_3$ $[M+H]^+$: 564.2849; found: 564.2855; $^1H$ NMR (500 MHz, $CDCl_3$) δ 1.29–1.54 (2H, m), 1.74–1.88 (2H, m), 1.90–2.12 (4H, m), 2.16 (3H, s), 2.33–2.62 (2H, m), 2.69 (3H, s), 2.78–2.94 (1H, m), 2.97 (3H, s), 3.05–3.31 (3H, m), 3.43-3.46 (1H, m), 3.51–3.65 (1H, m), 4.20–4.79 (5H, m), 5.99 (1H, br m), 7.11–7.48 (5H, m), 7.66–7.75 (1H, m); $^{13}C$ NMR (125 MHz, $CDCl_3$) δ 21.3, 21.8, 25.6, 25.7, 26.3, 27.8, 29.2, 29.8, 33.8, 35.8, 39.5, 39.7, 45.1, 49.6, 50.6, 52.9, 54.1, 56.9, 70.4, 71.4, 72.3, 85.8, 87.2, 123.0 - 130.0 (m), 136.6 - 140.5 (m), 169.6, 170.1, 170.8, 171.0; $^{19}F$ NMR (471 MHz, $CDCl_3$) δ -61.1 (3F), -204.7 (1F).

**2-(((3R,4S)-3-fluoropiperidin-4-yl)methoxy)-N-methyl-N-((S)-7-(3-((N-methylacetamido)methyl)phenyl)-5-(trifluoromethyl)-1,2,3,4-tetrahydronaphthalen-1-yl)acetamide (Compound A-(*S*))**

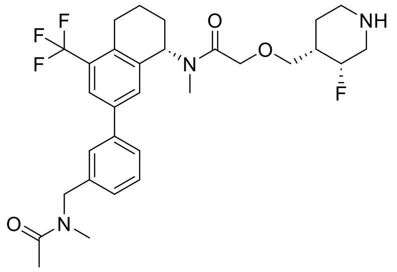

Prepared in an analogous method to **Compound A, Scheme 1,** where (*S*)-N-methyl-N-(3-(8-(methylamino)-4-(trifluoromethyl)-5,6,7,8-tetrahydronaphthalen-2-yl)benzyl)acetamide was used in place of *Rac*-N-methyl-N-(3-(8-(methylamino)-4-(trifluoromethyl)-5,6,7,8-tetrahydronaphthalen-2-yl)benzyl)acetamide. Resulting in the title compound (180 mg, 42 %) as an off-white solid.

Chiral HPLC, Column: CHIRALCEL OD-3, Mobile Phase: Hexane(0.1%DEA):EtOH=85:15, dr = >99.9:0.0; MS (ESI) *m/z* calcd for $C_{30}H_{37}F_4N_3O_3$ $[M+H]^+$: 564.3; found: 564.2. HRMS (ESI) *m/z* calcd for $C_{30}H_{37}F_4N_3O_3$ $[M+H]^+$: 564.2849; found: 564.2858; $^1H$ NMR (500 MHz, $CDCl_3$) δ 1.31–1.46 (2H, m), 1.75-1.82 (2H, m), 1.89–2.11 (4H, m), 2.14 (3H, s), 2.48-2.66 (2H, m), 2.68 (3H, s), 2.78–2.89 (1H, m), 2.94 (3H, s), 3.00–3.14 (2H, m), 3.16–3.25 (1H, m), 3.36–3.55 (2H, m), 4.18–4.33 (2H, m), 4.45–4.73 (3H, m), 6.02 (1H, br m), 7.20–7.47 (5H, m), 7.67-7.72 (1H, m); $^{13}C$ NMR (125 MHz, $CDCl_3$) δ 21.3, 21.8, 24.5, 25.7, 26.3, 27.9, 29.2, 29.9, 33.8, 35.7, 39.6, 39.9, 45.2, 49.6, 49.8, 50.6, 52.9, 54.1, 56.8, 70.4, 71.3, 72.2, 85.3, 87.6, 123.0 - 129.6 (m), 136.6 - 140.4 (m), 169.6, 170.1, 170.8, 171.0; $^{19}F$ NMR (471 MHz, $CDCl_3$) δ -61.1 (3F), -204.7 (1F).

**Synthesis of Compound B**

**Scheme 3**

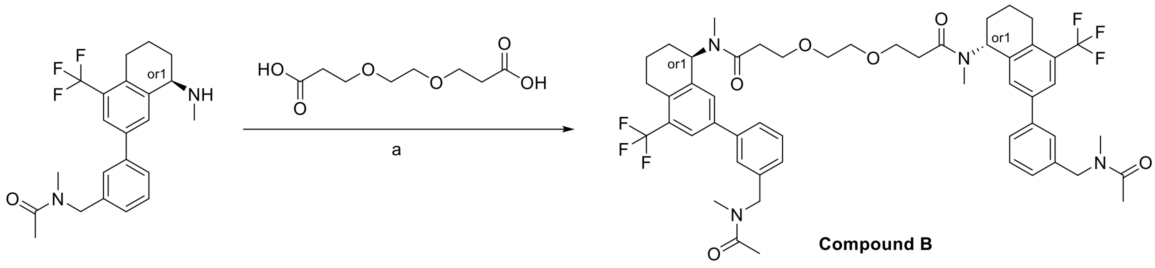

Reagents and conditions: (a) HATU, DIPEA, DMF, rt, 16 h

**3,3'-(ethane-1,2-diylbis(oxy))bis(N-methyl-N-(7-(3-((N-methylacetamido)methyl)phenyl)-5-(trifluoromethyl)-1,2,3,4-tetrahydronaphthalen-1-yl)propanamide) (Compound B)**

N-methyl-N-(3-(8-(methylamino)-4-(trifluoromethyl)-5,6,7,8-tetrahydronaphthalen-2-yl)benzyl)acetamide (49.2 mg, 0.13 mmol) was added to 3,3'-(ethane-1,2-diylbis(oxy))dipropionic acid (13 mg, 0.06 mmol), HATU (71.9 mg, 0.19 mmol) and DIPEA (0.066 mL, 0.38 mmol) in DMF (2 mL). The resulting mixture was stirred at rt for 16 hours. The reaction mixture was quenched with water (20 mL), extracted with EtOAc (3 x 10 mL), the organic layer was dried over $Na_2SO_4$, filtered and evaporated to afford white solid. The crude product was purified by flash C18-flash chromatography, elution gradient 0 to 80% MeCN in water (0.1%NH4HCO3). Pure fractions were evaporated to dryness to afford 3,3'-(ethane-1,2-diylbis(oxy))bis(N-methyl-N-(7-(3-((N-methylacetamido)methyl)phenyl)-5-(trifluoromethyl)-1,2,3,4-tetrahydronaphthalen-1-yl)propanamide) (30.0 mg, 50 %) as a yellow solid.

MS (ESI) *m/z* calcd for $C_{52}H_{60}F_6N_4O_6$ $[M+H]^+$: 951.4; found: 951.4. HRMS (ESI) *m/z* calcd for $C_{52}H_{60}F_6N_4O_6$ $[M+H]^+$: 951.4495; found: 951.4514; $^1$H NMR (400 MHz, DMSO-*d6*) δ 1.78–1.88 (m, 5H), 1.90-2.08 (m, 9H), 2.53-3.99 (m, 16H), 3.35 (s, 4H), 3.44 (s, 4H), 3.59-3.75 (m, 4H), 4.43–4.79 (m, 4H), 5.04–5.86 (m, 2H), 7.18-7.26 (m, 2H), 7.34–7.62 (m, 8H), 7.63–7.95 (m, 2H); $^{19}$F NMR (377 MHz, DMSO-*d6*) δ -59.8 (6F).

**Vibrational Circular Dichroism (VCD)**

**Experimental VCD:** The samples **4a** (Isomer 1) and 4b (Isomer 2) were both dissolved in $CDCl_3$ at a concentration of 0.184 M. The solutions were separately transferred to a 0.0995 mm $BaF_2$ cell and VCD spectra acquired for eight hours in a Biotools ChiralIR2X instrument at a resolution of 4 $cm^{-1}$ and PEM setting of 1400 $cm^{-1}$. A blank spectrum of the solvent was also acquired. Infra-red spectra were acquired on the samples concomitantly. The experimental infra-red spectra are shown in Figure S1 and experimental VCD spectra shown in Figure S2.

**Supplementary Fig. 1**. Experimental infra-red spectra for **4a** (red), **4b** (purple) and $CDCl_3$ solvent alone (blue).

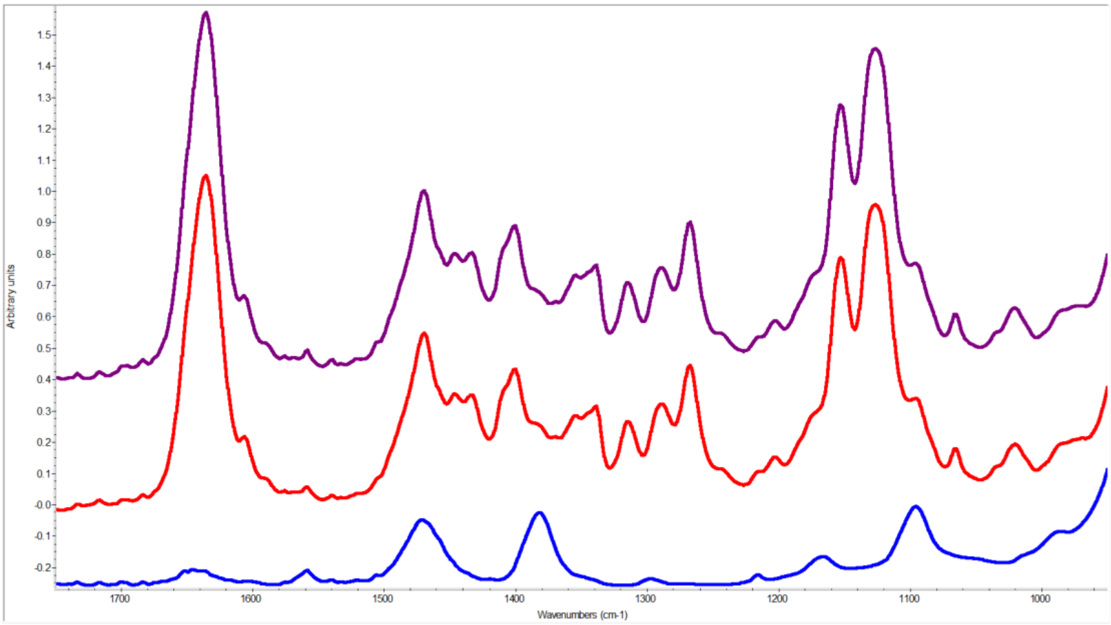

**Supplementary Fig. 2**. Experimental VCD spectra for **4a** (red), **4b** (purple) and $CDCl_3$ solvent alone (blue).

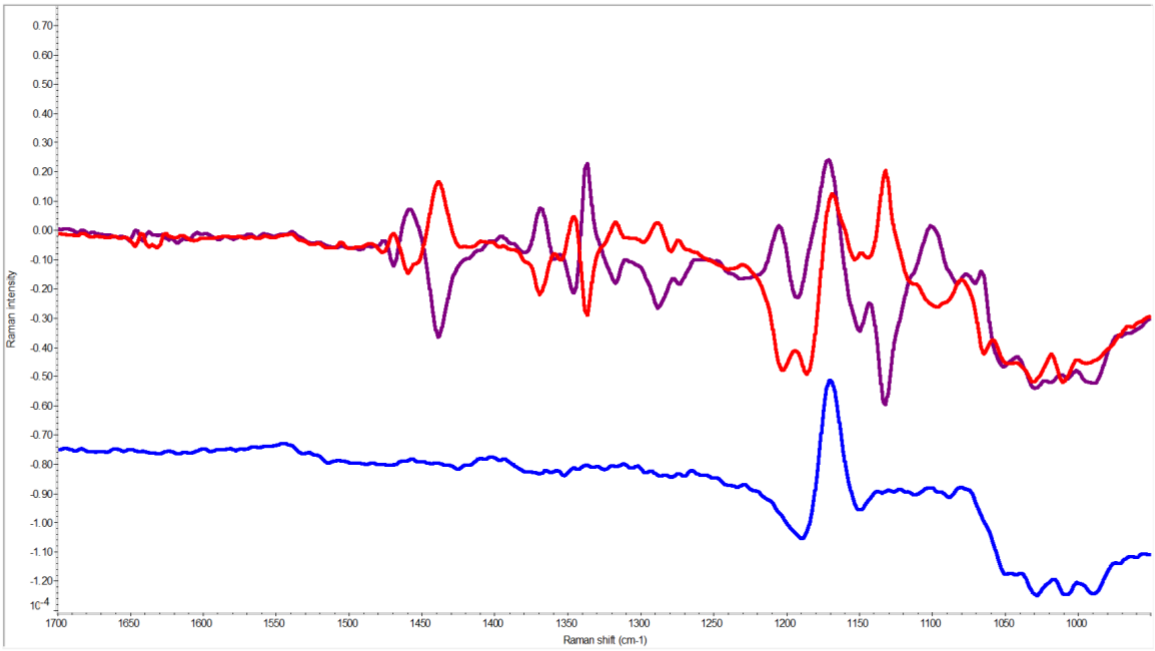

**Computational Spectral Simulations:** A Monte Carlo molecular mechanics search for low energy geometries was conducted for the *R* enantiomer. *MacroModel* within the *Maestro* graphical interface (Schrödinger Inc.) was used to generate 227 starting coordinates for conformers within 21 $kJmol^{-1}$ of the lowest energy conformer. These were used as starting points for density functional theory (DFT) minimizations within *Gaussian16*.[REF] Optimized structures, harmonic vibrational frequencies/intensities, VCD rotational strengths, and free energies at STP (including zero-point energies) were determined at the B3PW91/cc-pVTZ level of theory in the gas phase. After minimization, 41 conformations were found within 5 $kJmol^{-1}$ of the minimum and these are shown overlaid in Figure S3. The coordinates of the minimum energy conformation are shown in Table S1.

**Supplementary Fig. 3**. Overlay of the 41 lowest energy conformations (within 5 kJmol$^{-1}$ of the minimum) used in the calculation of the Boltzmann average IR and VCD spectra.

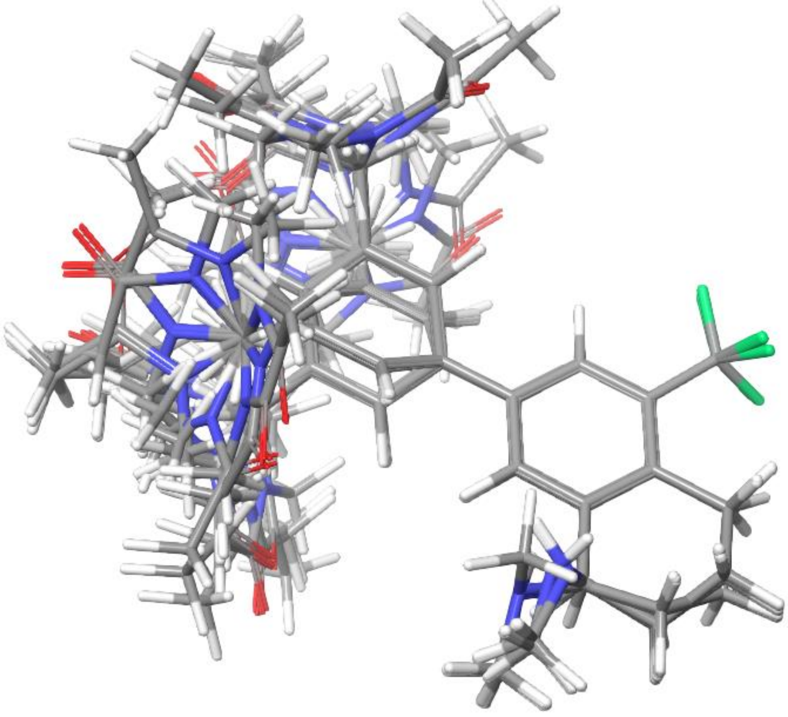

**Supplementary Table 2.** Coordinates for the minimum energy conformation.

| | | | |
|---|---|---|---|
| C | -2.15540 | -1.04110 | 0.03430 |
| C | -0.97080 | -1.51640 | -0.53750 |
| C | -1.00410 | -2.73520 | -1.21350 |
| C | -2.17870 | -3.47360 | -1.32660 |
| C | -3.34370 | -2.97920 | -0.75120 |
| C | -3.32890 | -1.76750 | -0.07170 |
| H | -2.14920 | -0.10950 | 0.58650 |
| H | -0.10180 | -3.12000 | -1.67400 |
| C | -2.16880 | -4.79590 | -2.05630 |
| H | -4.26570 | -3.54460 | -0.83110 |
| H | -4.23760 | -1.39090 | 0.38200 |
| C | 2.71540 | 0.71270 | -0.21470 |
| C | 1.47480 | 1.35720 | -0.36820 |
| C | 0.29290 | 0.64160 | -0.46320 |
| C | 0.28830 | -0.75180 | -0.42150 |
| C | 1.51480 | -1.38760 | -0.27510 |
| C | 2.71170 | -0.68510 | -0.16710 |
| C | 4.01270 | 1.48290 | -0.14940 |
| C | 1.39200 | 2.86160 | -0.43200 |

| | | | |
|---|---|---|---|
| H | -0.63680 | 1.17470 | -0.60080 |
| H | 1.54810 | -2.47140 | -0.23210 |
| C | 3.97720 | -1.49600 | 0.03080 |
| H | 3.91330 | -2.36200 | -0.64950 |
| C | 5.22540 | -0.69270 | -0.32420 |
| N | 4.03930 | -1.95150 | 1.42180 |
| H | 6.11610 | -1.25440 | -0.03580 |
| H | 5.27340 | -0.56410 | -1.41070 |
| C | 5.18370 | 0.65900 | 0.36560 |
| H | 6.11270 | 1.21030 | 0.20540 |
| H | 5.08370 | 0.49610 | 1.44130 |
| H | 3.88660 | 2.37730 | 0.46050 |
| H | 4.24140 | 1.84300 | -1.15860 |
| C | 4.82890 | -3.15140 | 1.62550 |
| H | 3.10220 | -2.09050 | 1.77390 |
| H | 4.74400 | -3.46720 | 2.66600 |
| H | 4.53130 | -3.99450 | 0.98190 |
| H | 5.88520 | -2.95000 | 1.43670 |
| F | 1.77150 | 3.44160 | 0.72590 |
| F | 2.17470 | 3.37910 | -1.39830 |
| F | 0.14560 | 3.29680 | -0.68000 |
| H | -3.17230 | -5.23110 | -2.03950 |
| N | -1.73120 | -4.69870 | -3.44060 |
| H | -1.48660 | -5.48870 | -1.56420 |
| C | -0.48440 | -5.14790 | -3.77160 |
| C | -2.68070 | -4.12180 | -4.36400 |
| O | 0.28860 | -5.57920 | -2.92910 |
| C | -0.08180 | -5.09520 | -5.22980 |
| H | -3.01890 | -3.14890 | -3.99770 |
| H | -2.23580 | -3.97430 | -5.34260 |
| H | -3.55970 | -4.76570 | -4.47780 |
| H | -0.78400 | -5.63060 | -5.87070 |
| H | -0.01610 | -4.06620 | -5.58930 |
| H | 0.89850 | -5.55610 | -5.30980 |

**Fit between calculated and experimental IR and VCD spectra:** An in-house program was used to generate a Boltzmann weighted average spectrum for the 41 conformations with the 5 $kJmol^{-1}$ limit and to fit Lorentzian line shapes (12 $cm^{-1}$ line width) to the computed spectra applying a linear scaling factor of 0.98. The fits between calculated and experimental infra-red and VCD data are shown in Figures S4 and S5 respectively.

There is very good agreement between calculated and experimental infra-red spectra, with only small differences between calculated and experimental frequencies noted for a few peaks. There is also good agreement between the calculated VCD spectrum for the (*R*) enantiomer and the experimental spectrum for **4a** with nearly every calculated peak agreeing in sign with the experimental spectrum. This is therefore very strong evidence that **4a** is the (*R*) enantiomer.

**Supplementary Fig. 4**. Comparison of calculated and experimental infra-red spectra. Calculated data for the (*R*) enantiomer is shown in green and experimental spectra for **4a** in red and **4b** in purple. The blank spectra recorded for $CDCl_3$ have been subtracted from both experimental spectra.

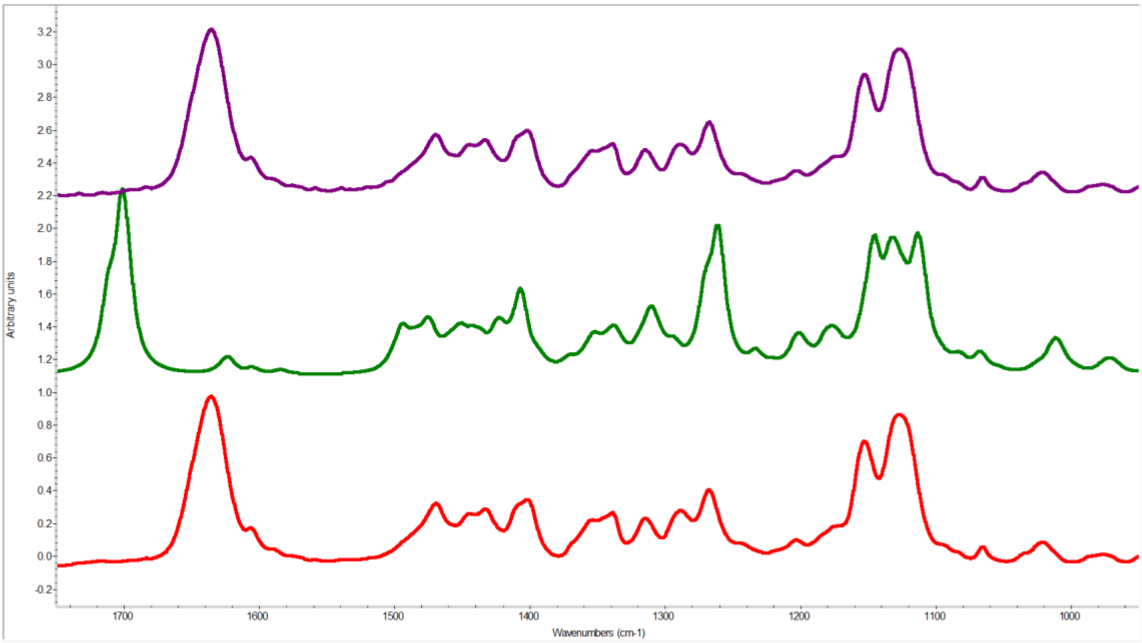

**Supplementary Fig. 5**. Comparison of calculated and experimental VCD spectra. Calculated data for the (*R*) enantiomer is shown in green and experimental spectra for **4a** in red and **4b** in purple. The blank spectra recorded for $CDCl_3$ have been subtracted from both experimental spectra.

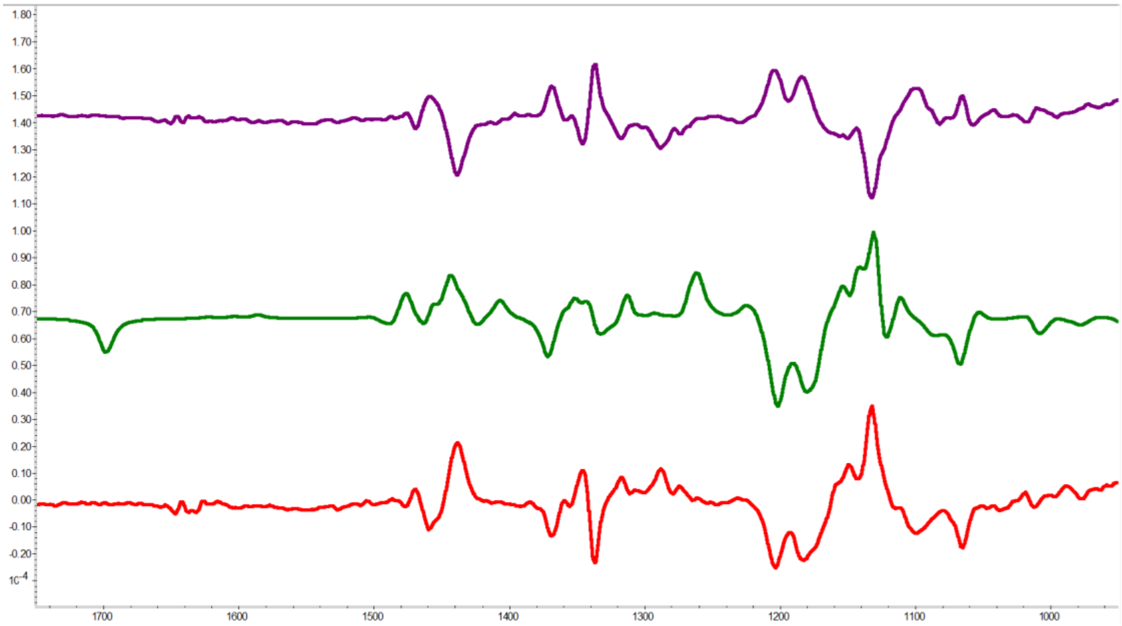

### References

Gaussian 16, Revision B.01, M. J. Frisch, G. W. Trucks, H. B. Schlegel, G. E. Scuseria, M. A. Robb, J. R. Cheeseman, G. Scalmani, V. Barone, G. A. Petersson, H. Nakatsuji, X. Li, M. Caricato, A. V. Marenich, J. Bloino, B. G. Janesko, R. Gomperts, B. Mennucci, H. P. Hratchian, J. V. Ortiz, A. F. Izmaylov, J. L. Sonnenberg, D. Williams-Young, F. Ding, F. Lipparini, F. Egidi, J. Goings, B. Peng, A. Petrone,

T. Henderson, D. Ranasinghe, V. G. Zakrzewski, J. Gao, N. Rega, G. Zheng, W. Liang, M. Hada, M. Ehara, K. Toyota, R. Fukuda, J. Hasegawa, M. Ishida, T. Nakajima, Y. Honda, O. Kitao, H. Nakai,

T. Vreven, K. Throssell, J. A. Montgomery, Jr., J. E. Peralta, F. Ogliaro, M. J. Bearpark, J. J. Heyd, E. N. Brothers, K. N. Kudin, V. N. Staroverov, T. A. Keith, R. Kobayashi, J. Normand, K. Raghavachari, A. P. Rendell, J. C. Burant, S. S. Lyengar, J. Tomasi, M. Cossi, J. M. Millam, M. Klene, C. Adamo, R. Cammi, J. W. Ochterski, R. L. Martin, K. Morokuma, O. Farkas, J. B. Foresman, and D. J. Fox, Gaussian, Inc., Wallingford CT, 2016.
